# Discovery and Validation of Alternatives to VSV-G for Pseudotyping of Lentiviral Vectors for In Vivo Delivery of Anti-Tumor Transgenes

**DOI:** 10.1101/2025.03.03.641199

**Authors:** M.J. Spindler, A. Amezquita, E.F.X. Byrne, R. Edgar, S. Ravi, S. Sandhu, T. Weller, D.S. Johnson

## Abstract

Though cell therapy for cancer is now widely used commercially and has been efficacious for tens of thousands of patients, conventional manufacturing methods are expensive and difficult to scale. An alternative approach is to deliver the relevant anti-tumor transgenes to T cells *in vivo* in the patient. Such “*in vivo* cell therapy” methods promise to be more scalable, with reduced cost of goods, since the same drug product can be administered to any patient. Typically, conventional cell therapy introduces anti-tumor transgenes into T cells using a lentivector pseudotyped with VSV-G. However, VSV-G is not cell type specific because its molecular target is present on a diversity of human cells, which may result in less than optimal pharmacology *in vivo*. Nature’s existing viral diversity presents the opportunity to identify alternative pseudotypes that are more optimal for *in vivo* cell therapy. In this study, we first performed a large-scale bioinformatic sequence search for G-proteins similar to VSV-G. We identified 166 G-proteins in the sequence search and then tested 9 *in vitro* for efficiency, specificity, and sensitivity of transgene delivery to various human immune cell phenotypes. We used the results from this screen to select three G-protein candidates for a pilot GFP transgene delivery study in humanized mice, using anti-CD3 antibody fragments for T cell tropism. One candidate G-protein performed significantly better than VSV-G, so we moved that candidate into tumor control studies in a humanized mouse model. This candidate G-protein was able to deliver an anti-tumor transgene to T cells, which subsequently cleared 100% of tumor burden in 100% of mice. We conclude that systematic screens for optimal lentivector designs can be used to identify optimized candidates for *in vivo* cell therapy.

## BACKGROUND

Adoptive cell therapies wherein chimeric antigen receptors (CARs) are engineered into autologous T cells *ex vivo* have shown strong clinical efficacy and safety, with four anti-CD19 CAR-T cell therapies (Breyanzi, Kymriah, Tecartus, and Yescarta) and two anti-B cell maturation antigen (BCMA) CAR-T cell therapies (Abecma and Carvykti) now approved by FDA.

Despite these transformative successes in the clinic, many barriers, including treatment costs and limited manufacturing slots, continue to restrict patient access to these lifesaving therapies (Mikhael et al., 2022). On average, the cost of the engineered CAR-T cell medication is about $530,000, and the total cost of care typically exceeds $1 million (Antrim, 2021). Manufacturers cite the expensive, labor-intensive, and personalized manufacturing process, which totals $150,000-$300,000 per patient (Vormittag et al., 2018). Indeed, CAR-T manufacturing is complex, requiring the manufacturing of CAR-T lentivirus and patient specific peripheral blood mononuclear cell (PBMC) collection, T cell isolation, *ex vivo* T cell stimulation, transduction, and expansion. In addition, there are significant requirements for controls and characterization of the CAR-T product prior to infusion.

To decrease costs and increase production scalability, CAR-T cell manufacturers have focused on operational efficiencies. Personnel costs are as high as 72% of manufacturing costs with current approaches (Spink & Steinsapir, 2018; Ran et al., 2020), highlighting the potential of roboticization to reduce costs (Winn, 2022). Other product developers have focused on scaling manufacturing by developing off-the-shelf allogeneic cell therapy approaches, for example, CAR-NK cells, CAR-macrophage cells, and human leukocyte antigen (HLA)-deficient and/or T cell receptor (TCR)-deficient CAR-T cells (e.g., Pan et al., 2022). These methods have the advantage of scalability to thousands of doses per manufacturing run but remain largely unproven clinically.

An innovative alternative approach is to engineer T cells *in vivo* using gene delivery vectors that specifically target T cell populations. *In vivo* delivery of CAR genes to T cells could significantly reduce manufacturing costs, and time to treatment, by removing *ex vivo* cell handling altogether. Lentivirus is the most common method for engineering CAR-T cells *ex vivo* and is widely considered safe in that context (Poletti & Mavilio 2021), but the conventional lentiviral pseudotype, VSV-G, targets the broadly expressed low density lipoprotein receptor (LDL-R), which could result in low efficiency and specificity when delivered *in vivo*.

Starting about two decades ago, investigators have added antibody fragments (single chain variable fragments, or scFvs) to VSV-G pseudotyped lentivectors, to direct cell type-specific tropism, with the goal of *in vivo* gene delivery (Yang et al., 2006). Other groups have suggested improved specificity by combining scFvs with a mutant VSV-G (“mut-VSV-G”) that disrupts native LDL-R binding (Yu et al., 2022; Dobson et al., 2022). In another study, lentivectors loaded with anti-CD3 scFvs and pseudotyped with the Cocal virus envelope glycoprotein, delivering an anti-CD19 CAR, achieved clearance of aggressive B cell lymphoma tumors in a mouse model (Michels et al., 2023). Such methods show promise for *in vivo* delivery of CAR-T, drastically reducing the cost of cell therapy while retaining efficacy and safety but require further optimization.

Despite the enormous opportunities available in the field of *in vivo* transgene delivery with lentivectors, only a handful of pseudotypes have been tested preclinically. In this study, we explore alternative pseudotypes to VSV-G, mut-VSV-G, and Cocal virus, which may eventually be shown to have advantages in terms of efficiency, specificity, sensitivity, immunogenicity, stability, and/or manufacturability. First, we did a bioinformatic sequence search of public sequencing databases for viral envelope glycoproteins (G-proteins) that are structurally related to VSV-G. We chose nine G-proteins with sequence identities to VSV-G from 26% to 85% and tested these *in vitro* using human PBMCs and flow cytometry. Next, we tested three pseudotypes in a pilot mouse study showing efficiency of gene delivery. Finally, we tested one candidate pseudotype in a NALM6 B cell acute lymphoblastic leukemia (ALL) xenograft mouse model and showed 100% tumor clearance in 100% of mice.

## MATERIALS AND METHODS

### Sequence Search and Analysis

First, we used Serratus (Edgar et al., 2022), a biological sequence search engine which is highly optimized for Amazon Web Services (AWS) infrastructure, to search for proteins with similarity to VSV-G (Genbank QEL51172.1). Serratus was run using the diamond amino acid alignment module (https://github.com/ababaian/serratus/tree/master/containers/serratus-align) to search the NCBI non-redundant protein database (NR) and the Transcriptome Shotgun Assembly (TSA) sequencing database in GenBank (https://www.ncbi.nlm.nih.gov/genbank/tsa/). This search identified 166 unique G-proteins with protein sequence similarity to VSV-G. We used CLUSTAL O (1.2.4) to generate a multiple sequence alignment (MSA) and phylogenetic tree (**Supplementary Figures S1-S2**) for these 166 proteins and then selected 9 representative G-proteins (“Candidates” A through I) for further study. CLUSTAL O was again used to generate an MSA, amino acid identities, and a putative phylogenetic tree for the 9 G-proteins. We generated putative protein structures for the 9 representative G-proteins with ColabFold (an implementation of AlphaFold2) on Google Colab (Mirdita et al., 2022). For all proteins, we used a custom template (VSV-G prefusion experimental structure PDB 5i2s) to induce a “prefusion” conformation. For the purpose of coloring amino acids conserved with VSV-G, each of Candidates A through I were aligned to the putative VSV-G structure using the PyMol Align tool.

### Adherent Lentivector Packaging for In Vitro Studies

Third generation lentivirus was packaged using equimolar amounts of packaging plasmids: pRSV-Rev (Cell Biolabs), pCgpV (Cell Biolabs) or pLP1 (Thermo Fisher), pTwist-CMV-pseudotype (synthesized at Twist Bio), mRuby, GFP, or anti-CD19 CAR-GFP lentiviral transfer plasmids (synthesized in-house using gBlocks and Gibson Assembly, and sequence verified), and appropriate cell targeting scFv constructs (synthesized in-house using gBlocks and Gibson Assembly, and sequence verified) in Lenti-Pac 293Ta cells (GeneCopoeia). The scFvs were tethered to an extracellular Fc domain and a CD28 transmembrane domain (lacking the intracellular signaling domain). Plasmids were reverse transfected into Lenti-Pac 293Ta cells using Lipofectamine 3000 (ThermoFisher) and lentiviral supernatant was harvested at 24 and 48 hours post transfection and pooled. Lentiviral supernatant was centrifuged at 2000×g for 10 minutes to remove cellular debris and filtered through a 0.45μm filter. As needed, lentivector preparations were concentrated using Lenti-X Concentrator (Takara) and pelleted viral particles were resuspended in cell culture media for an about 20× concentration. Lentivector particle (LP) counts were determined using viral RNA isolated with the NucleoSpin RNA Virus Kit (Macherey-Nagel) and the Lenti-X qRT-PCR Titration Kit (Takara).

### Lentivector Screening in Human PBMCs

In pilot studies, cryopreserved human PBMCs were thawed into human PBMC culture media (IMDM media containing 5% human AB serum, 100 IU/mL IL-2, and Penicillin-Streptomycin) and seeded at a cell density of 2×10^6^ cells/mL for overnight recovery. The following day, human PBMCs were seeded for a final cell density of 1×10^6^ cells/mL and incubated with freshly concentrated lentivector in the presence of 8μg/mL Polybrene (Santa Cruz Biotechnology) for 2 days. After incubation, lentivector transduction was measured by flow cytometry. PBMC samples were stained for surface CD3, CD4, CD8A, CD19, and CD20 expression and assessed for cell viability using DAPI staining and mRuby expression. Flow cytometry data was analyzed in FlowJo (BD) (**Supplementary Figure S3**). Subsequent studies found that the addition of Polybrene did not improve transduction of human PBMCs when cell targeting moieties were present on the lentivector.

In the pseudotype by cell targeting moiety combination screen, cryopreserved human PBMCs were thawed into human PBMC culture media and seeded at a cell density of 2×10^6^ cells/mL for overnight recovery. The following day, 5×10^5^ human PBMCs were seeded for a final cell density of 1×10^6^ cells/mL. Cryopreserved lentivectors were thawed and diluted in human PBMC culture media to normalize the number of lentivector particles (LPs) within each set of targeting moieties. LP counts were determined using the Lenti-X qRT-PCR Titration Kit as described above. LP doses were: No Targeting at 2.43×10^8^ LPs; CD3 Targeting at 1.66×10^8^ LPs; CD4 Targeting at 1.51×10^8^ LPs; CD8 Targeting at 1.78×10^8^ LPs (exceptions for anti-CD8 due to lower packaging yields: no pseudotype at 1.12×10^8^ LPs and Candidate D at 1.11×10^8^ LPs); and CD19 Targeting at 1.57×10^8^ LPs.

Human PBMCs were incubated with lentivector for 3 days and then analyzed for transduction by flow cytometry. PBMC samples were stained for surface CD3, CD4, CD8A, CD19, CD20, and TCRab expression and assessed for cell viability using DAPI staining and GFP expression. Flow cytometry data was analyzed in FlowJo (**Supplementary Figure S4**). We used the processed flow cytometry data to calculate the efficiency, sensitivity, and specificity of each lentivector sample. The efficiency calculation allows us to compare transduction levels across the different cell targeting moieties by normalizing for the input LP count.

Calculations:

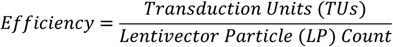

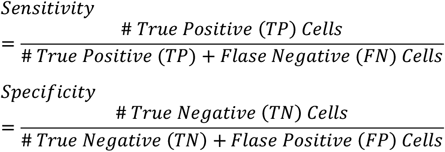

Where transduction units (TUs) were determined by multiplying the frequency (%) of transduced (GFP^+^) on-target cells by the estimated number of on-target cells in the starting PBMC population. The estimated number of on-target cells was determined from no virus treatment samples: CD3 Targeting: 339,928 cells based on live / CD4^+^ and/or CD8^+^ gating; CD4 Targeting: 209,419 cells based on live / CD3^+^ / CD8^-^ gating; CD8 Targeting: 128,166 cells based on live / CD3^+^ / CD4^-^ gating; CD19 Targeting: 71,250 cells based on live / CD20^+^ gating. True positive (TP) cells are transduced (GFP^+^) surface marker positive cells. False negative (FN) cells are non-transduced (GFP^-^) surface marker positive cells. True negative (TN) cells are non-transduced (GFP^-^) surface marker negative cells. False positive (FP) cells are transduced (GFP^+^) surface marker negative cells. Surface marker positive and negative cell populations are shown in **Supplementary Table S1**.

Antibodies used for flow cytometry are listed in **Supplementary Table S2**.

### Generation NALM6 Cells of Luciferase-Expressing

A multicistronic lentiviral transfer construct encoding firefly luciferase (ffLuc)-T2A-mRuby3-P2A-Puromycin resistance genes driven by the EF1alpha promoter was transduced into NALM6 cells (ATCC, CRL-3273). T2A and P2A are ribosome skip motifs derived from thosea asigna virus 2A and porcine teschovirus-1 2A, respectively. A population of stably integrated ffLuc-mRuby3-PuroR cells were enriched by selection in media containing 0.5 ug/mL Puromycin for 2 weeks followed by two successive rounds of cell sorting for mRuby3+ cells. Cells were cultured in media without Puromycin selection after the initial 2 week selection process. Stable mRuby3 expression was validated by flow cytometry (see **Supplementary Figure S5A)**. Firefly luciferase activity was validated with *in vitro* live cell bioluminescent assays over 7 weeks in culture (see **Supplementary Figure S5B)**. Briefly, 5×10^5^ cells were pelleted and resuspended in 0.5mL media containing 150 μg/mL IVISbrite D-Luciferin Potassium Salt Bioluminescent Substrate (Revvity). Cells were seeded at 0.1mL into quadruplicate wells of a white 96-well assay plate and incubated at 37ºC for 10 minutes. Bioluminescence signal was read on a Promega GloMax system.

### Suspension Lentivector Packaging for In Vivo Studies

Third generation lentivirus was packaged using the LV-MAX Lentiviral Production System (Thermo Fisher) following the manufacturers recommendations. Briefly, 2.4×10^8^ cells were transient transfected with 36μg pRSV-Rev (Cell Biolabs), 18μg pLP1 (Thermo Fisher), 36μg of the appropriate viral envelope plasmid (i.e., pseudotype), 18μg of the human CD3-targeting moiety plasmid, and 120μg of the appropriate lentiviral transfer plasmid (GFP or anti-CD19 CAR-GFP) in 60mL final production volume. Lentiviral supernatant was harvested at 48 hours post transfection, clarified by centrifugation at 1300×g for 15 minutes and filtration through 0.45μm filters, and treated with 15 U/mL Benzonase (Sigma-Aldrich) for 30 minutes at 37ºC. Lentivector was concentrated by high-speed centrifugation at 20,000×g for 2 hours at 4ºC using a 20% Sucrose cushion. Pelleted viral particles were resuspended in DPBS and cryopreserved at −80ºC until use. Viral RNA was isolated from freshly resuspended viral particles using the NucleoSpin RNA Virus Kit (Macherey-Nagel) and lentivector particle (LP) counts were determined using the Lenti-X qRT-PCR Titration Kit (Takara). Functional titers, transducing units per mL (TUs/mL), were determined by transducing 1×10^5^ Jurkat E6-1 (ATCC) cells with a dilution series of previously cryopreserved lentivector particles and assessing viable cells for GFP expression by flow cytometry after 3 days of viral incubation. The functional titration ED50 was calculated from these dose-responses using R package `drc` with the following model parameters: `model<-drm(response∼dose,fct=LL.2(), type=“Poisson”)`.

The R estimated ED50 value was used to calculate TUs/mL:

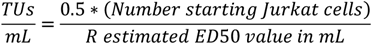

Lentivector particle (LP) count and transduction unit (TU) concentrations and yields for the *in vivo* mouse studies are provided in **Supplementary Table S3**.

### In Vivo *Mouse Studies*

Mouse studies used the Nod scid gamma Major Histocompatibility Complex double knockout strain (NOD.Cg-*Prkdc*^*scid*^ *H2-K1*^*b-tm1Bpe*^ *H2-Ab1*^*g7-em1Mvw*^ *H2-D1*^*b-tm1Bpe*^ *Il2rg*^*tm1Wjl*^/SzJ; Jackson Labs strain 025216); we will refer to this mouse strain as “NSG MHC DKO”. Mispro (South San Francisco, CA) performed routine vivarium services (including daily cage checks), veterinary care, and Institutional Animal Care and Use Committee (IACUC) review.

For the *in vivo* transduction efficiency pilot study, lentivectors expressing anti-CD3 scFv and pseudotyped with VSV-G, mut-VSV-G, or Candidates B, C, or I, were synthesized and characterized as above, using a GFP transgene construct. On Day 0, 7 week old, female, NSG MHC DKO mice were injected retro-orbitally (r.o.) into the right eye with 10^7^ human PBMCs. 45-60 minutes later, mice were injected r.o. into the left eye with 2×10^10^ LPs for one lentivector design (n=3 mice per lentivector design). On Day 8, 100μL of blood was drawn via cheek puncture, and flow cytometry was performed. Mouse whole blood was processed with anti-mouse CD16/CD32 (Mouse BD Fc Block; BD Pharmingen) and human FcR Blocking Reagent (Miltenyi Biotec), stained for surface human (h)CD45, hCD3, hCD4, hCD8A, hCD19, and hCD20 expression, processed with a red blood cell lysis solution (Miltenyi Biotec or BioLegend), and assessed by flow cytometry for cell viability using DAPI, GFP expression, and antibody staining. Flow cytometry data was analyzed in FlowJo (**Supplementary Figure S6A**).

For the mouse tumor control study, lentivectors expressing anti-human CD3 scFv and pseudotyped with Candidate C were synthesized and characterized as above, using either GFP only or anti-CD19 CAR-GFP transgene constructs. On Day −14 (D-14), 7 week old, female, NSG MHC DKO mice were injected with 10^6^ ffLuc-mRuby3 NALM6 cells intraperitoneally (i.p.). Whole-body bioluminescent imaging (BLI) was performed on Days −8 and −1 (D-8 and D-1) 10-20 minutes after IP injection of 150mg/kg IVISbrite D-Luciferin Potassium Salt Bioluminescent Substrate (Revvity) using a Pearl Trilogy Imaging System (LI-COR). BLI signal was quantified using Image Studio Software (v6.0; LI-COR) with background normalization to an empty region of the image. Mice were randomized by luminescence signal intensity on D-1. On Day 0 (D0), mice were either left untreated without PBMC or lentivector injections (Group 1, n=6) or injected r.o. into the right eye with 10^7^ human PBMCs (Group 2, n=7; and Group 3, n=7). 45-60 minutes later, 1.5×10^7^ TUs of GFP transgene lentivector (Group 2) or anti-CD19 CAR-GFP transgene lentivector (Group 3), were injected r.o. into the left eye. Mice were weighed and body conditions assessed twice a week for the duration of the study. Whole-body BLI were performed once a week through the end of the study.

Flow cytometry was performed once a week on 100μL of blood drawn via cheek puncture. Mouse whole blood was processed with anti-mouse CD16/CD32 (Mouse BD Fc Block; BD Pharmingen) and human FcR Blocking Reagent (Miltenyi Biotec), stained for surface mouse (m)CD45, human (h)CD45, hCD3, hCD4, hCD8A, hCD19, and hCD20 expression, processed with a red blood cell lysis solution (Miltenyi Biotec or BioLegend), and assessed by flow cytometry for cell viability using DAPI, GFP expression, and antibody staining. Flow cytometry data was analyzed in FlowJo (**Supplementary Figure S6B**). Antibodies used for flow cytometry are listed in **Supplementary Table S2**.

## RESULTS

### Bioinformatics identifies a set of viral G-proteins structurally similar to VSV-G

We set out to use bioinformatics to identify a small but diverse set of viral envelope G-proteins which share sequence and structural features with VSV-G (**Figure 1A**). Using the Serratus cloud-based algorithm (Edgar et al., 2022) to search public sequence databases, we identified 166 proteins with sequence similarity to VSV-G (**Supplementary Figures S1-S2**). This set of 166 G-proteins ranged in amino acid identity to VSV-G from 17.3% to 99.4%, with an average of 57.5% and a median of 49.1%. We arbitrarily chose nine G-proteins (pseudotyping Candidates A through I) with protein identities to VSV-G of 26% to 85% (average 40% and median 38% identity) for further *in silico* and *in vitro* analysis (**Figure 1B**). Of this set, only Candidate I (85% sequence identity to VSV-G) retains the three amino acid residues (H8, K47, R354) that when mutated abolish binding of VSV-G to its primary receptor target, LDL-R (Nikolic et al., 2018).

**Figure 1.**
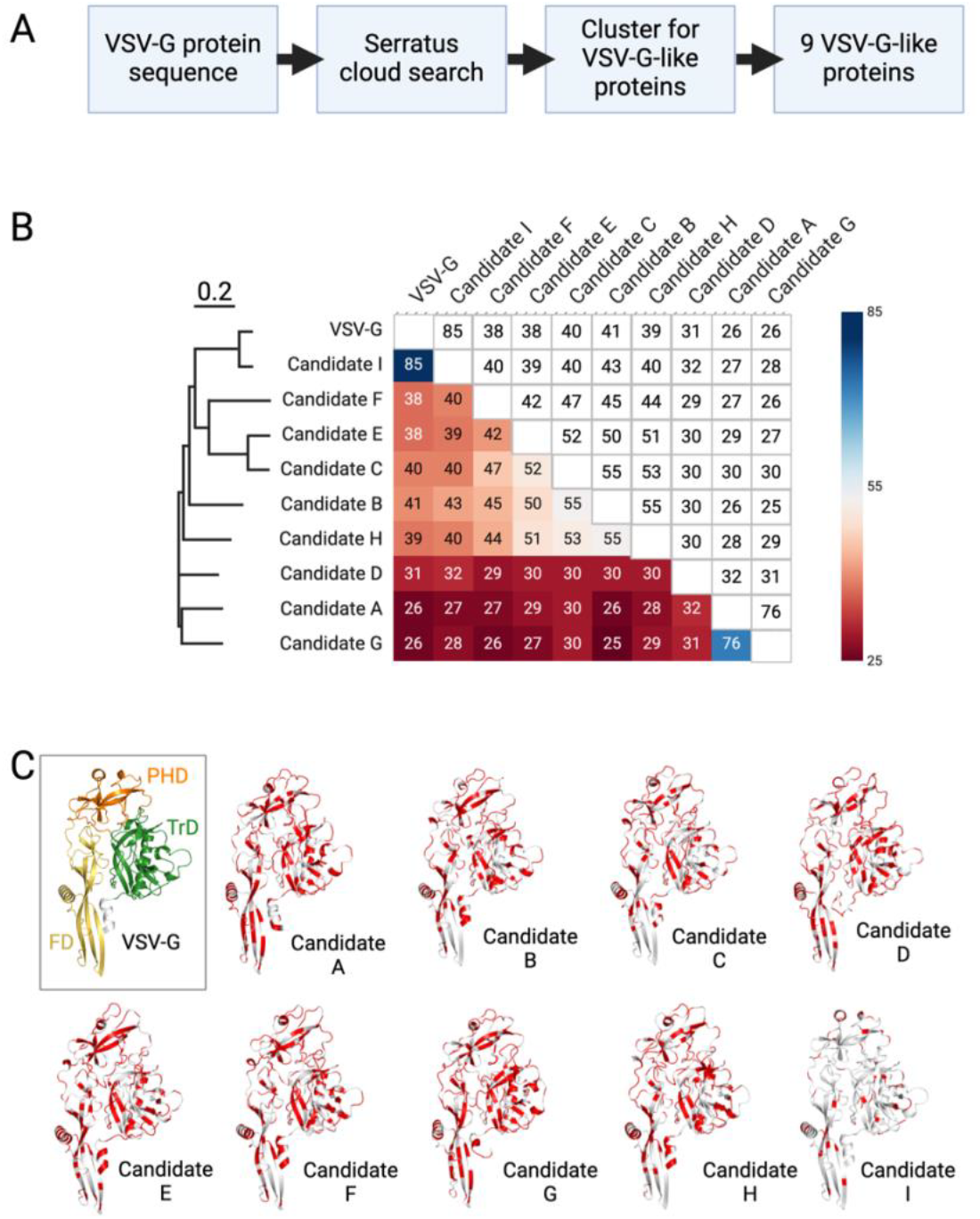
Discovery and bioinformatic characterization of candidate viral surface receptors for pseudotyping lentivectors for *in vivo* transgene delivery. (A) Bioinformatic protein sequence search workflow used to identify nine VSV-G-like protein candidates for pseudotyping. (B) Protein sequence identity matrix and tree for VSV-G and nine viral G-protein candidates. Blue to red color gradient represents percent protein sequence identity between a given pair of proteins. Numbers in the matrix (upper right half) indicate percent protein sequence identity between a given pair of proteins. Bar above the tree represents an evolutionary distance of 0.2 as computed by CLUSTAL O. (C) Predicted AlphaFold2 structures for VSV-G and nine G-protein candidates identified in the bioinformatic protein sequence search, shown in cartoon representation. For clarity, signal sequences, transmembrane and cytosolic regions of the predicted structures have been removed in the images (although they were included in the predictions). VSV-G is colored by domain: fusion domain (FD) in yellow, pleckstrin homology domain (PHD) in orange, trimerization domain (TrD) in green. Other structures are colored by sequence identity to VSV-G: identical residues in white, non-identical residues in red.

Next, we used artificial intelligence (AI) to predict protein structures for VSV-G and the nine selected VSV-G-like proteins (**Figure 1C**). VSV-G has multiple functions, including viral attachment to LDL-R, entry into the target cell via a clathrin-mediated endocytic pathway, and pH-dependent fusion of the viral envelope with the endosomal membrane. The VSV-G ectodomain folds into three distinct domains: the fusion domain (FD), the pleckstrin homology domain (PHD), and the trimerization domain (TrD). By visual inspection, the AI-predicted structures of all nine Candidates appear very similar to VSV-G, each containing all three domains with the same relative positioning. Indeed, upon structural alignment with VSV-G, each Candidate had root mean square deviations (rmsd) of Cα positions of just 0.25 – 2.1 Å. This very high structural similarity is observed despite some G-protein Candidates having amino acid sequence identities to VSV-G as low as 26%, reinforcing the utility of the Serratus search algorithm. The differences in sequence are distributed throughout the three domains. Aside from the LDL-R-binding triad of residues noted above, the functional effects of these differences are not readily predicted from the primary sequence or the predicted structures.

### High-throughput in vitro transduction of human PBMCs characterizes sensitivity, specificity, and efficiency of gene delivery using nine candidate viral G-proteins

Given the structural similarities in the context of high amino acid sequence divergence among Candidates A through I, we were interested in assessing the functional similarities and differences among the viral G-proteins. For example, a given G-protein may be more efficient at transduction of various cell types, or more specific for transduction of a particular cell type. To prepare for a cell-targeted viral G-protein screen, we ran a set of pilot studies testing (i) various pseudotyped lentivectors for their ability to transduce T cells and (ii) various cell-targeting moieties for their ability to direct lentivectors to specific cell types.

First, we tested lentivectors pseudotyped with VSV-G, mutant (mut)-VSV-G, or Candidate A (26% ID to VSV-G), B (41% ID to VSV-G), or C (40% ID to VSV-G) G-proteins with and without human CD3-targeting carrying an mRuby transgene for their ability to transduce human T cells within a heterogeneous PBMC population (**Figure 2A**). Non-targeting lentivector showed low to no transduction of human T cells whereas CD3-targeted lentivectors showed variable levels of T cell transduction. CD3-targeted lentivectors pseudotyped with VSV-G, mut-VSV-G and Candidate C all showed transduction efficiencies of 95% or greater, Candidate B had 60% and Candidate A had 42% transduction efficiencies. Additionally, CD3-targeted lentivectors induced blast formation in culture, indicating activation and expansion of the lentivector targeted T cells whereas non-targeted lentivector did not induce blast formation (data not shown). These results indicate that our set of highly sequence diverse viral G-proteins can mediate lentivector gene delivery to target cells with efficiencies similar to that of VSV-G.

**Figure 2.**
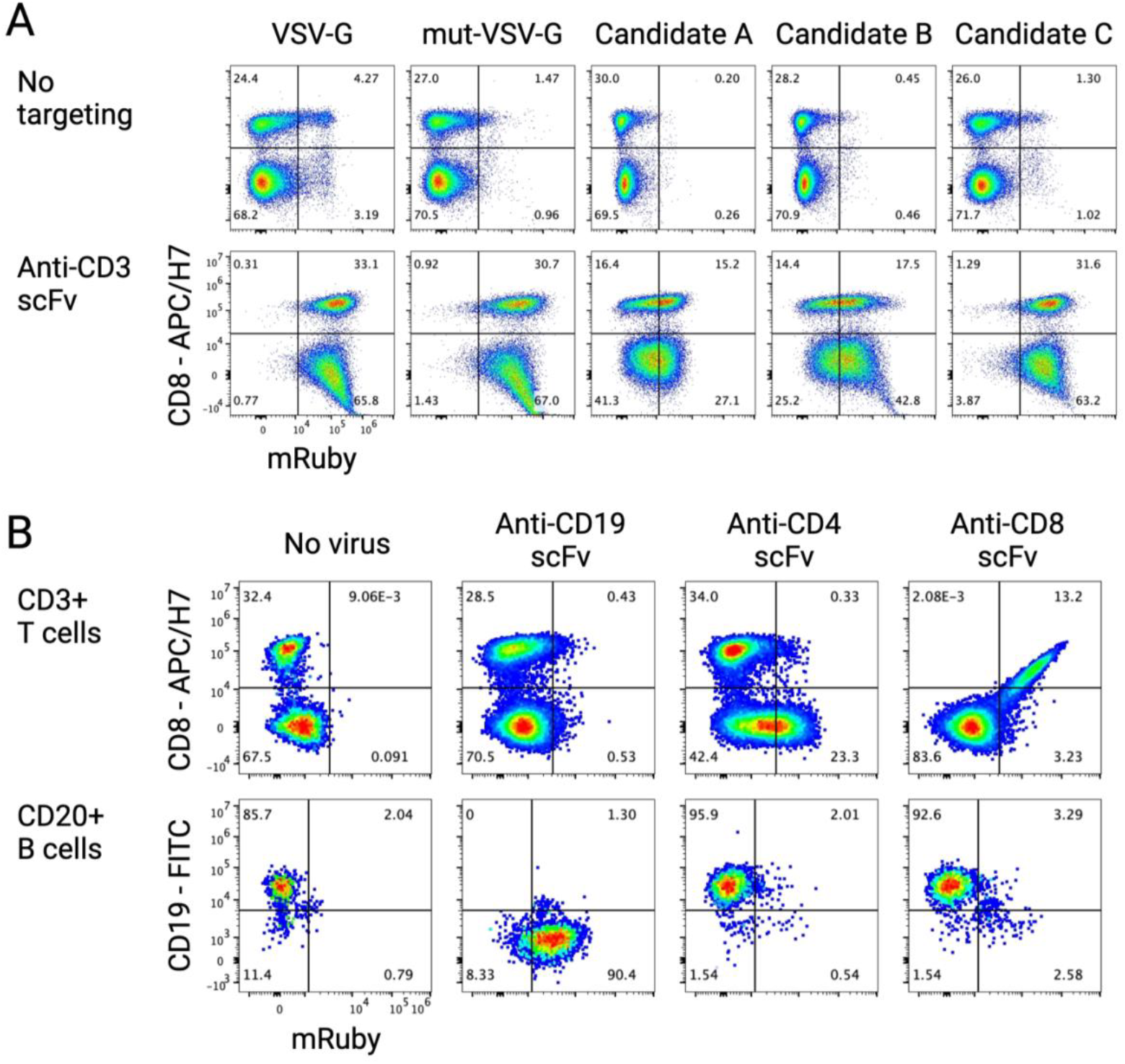
Pilot *in vitro* cell-targeted lentivector gene delivery studies in human PBMCs. (A) Flow cytometry data displaying CD8 surface expression versus mRuby fluorescent signal for PBMC samples transduced with non-targeted and CD3-targeted lentivectors pseudotyped with VSV-G, mut-VSV-G, Candidate A, B, or C. Data is from a parental gate for CD4 and/or CD8 positive cells. (B) Flow cytometry data displaying CD8 (top row) or CD19 (bottom row) surface expression versus mRuby fluorescent signal for PBMC samples transduced with Candidate C pseudotyped lentivector with anti-CD19, anti-CD4, or anti-CD8 scFv targeting. A no virus transduction negative control sample is also presented. Data is from parental gates for CD3 positive (top row) T cells or CD20 positive (bottom row) B cells.

Next, we tested lentivectors pseudotyped with Candidate C and cell targeting scFv moieties for CD19, CD4, and CD8 carrying an mRuby transgene for their ability to specifically transduce target cells in a heterogeneous human PBMC population (**Figure 2B**). The lentivectors all showed specific viral transduction of the targeted cell population. CD19-targeted lentivector transduced 90% of CD20^+^ B cells, CD4-targeted lentivector transduced 35% of CD8^-^ T cells, and CD8-targeted lentivector transduced 99% of CD8^+^ T cells. Unlike the CD3-targeted lentivectors, CD4 and CD8-targeted lentivectors did not induce blast formation in culture suggesting that these targeted lentivectors did not induce T cell activation (data not shown). These results indicate that candidate viral G-proteins can mediate gene delivery to various cell populations including non-activated T cells.

Following the successful completion of these pilot *in vitro* studies, we set up a larger-scale combinatorial screen to test all 9 candidate viral G-proteins side-by-side for targeted cell transduction. We packaged lentivector particles with viral G-protein Candidates A through I,VSV-G, or no viral G-protein, and either no scFv targeting, anti-CD3, anti-CD4, anti-CD8, or anti-CD19 scFv targeting. We used a lentiviral transfer construct with a GFP fluorescent reporter to mark the transduced cells. We transduced human PBMCs *in vitro* with a normalized dose of lentivector particles for each cell-targeting group and assessed GFP expression in various PBMC cell populations by flow cytometry (**Supplementary Figure S7**). Generally, the no-targeting lentivector samples show weak GFP expression levels in B and T cells (**Figure 3A**). We did not expect the non-targeted VSV-G pseudotyped lentivector to efficiently transduce B or T cells in this assay, as non-activated human B and T cells lack surface expression of LDL-R (Amarache 2014). As expected, the non-targeted VSV-G pseudotyped sample did not show a clear population of transduced B or T cells but just a slight shift in GFP signal compared to the non-targeted no pseudotype control with 2.7% of B cells and 0.92% of T cells showing weak GFP signal. In comparison, CD3-targeted VSV-G lentivector shows a clear population of high GFP^+^ T cells. The non-targeted candidate G-protein pseudotyped lentivectors showed similarly low levels of GFP signal in B and T cells with slightly higher frequencies of GFP^+^ cells for Candidates E (B cells: 4.1%; T cells: 2.9%) and C (B cells: 16.6%; T cells: 5.1%) but again no clear population of transduced cells.

**Figure 3.**
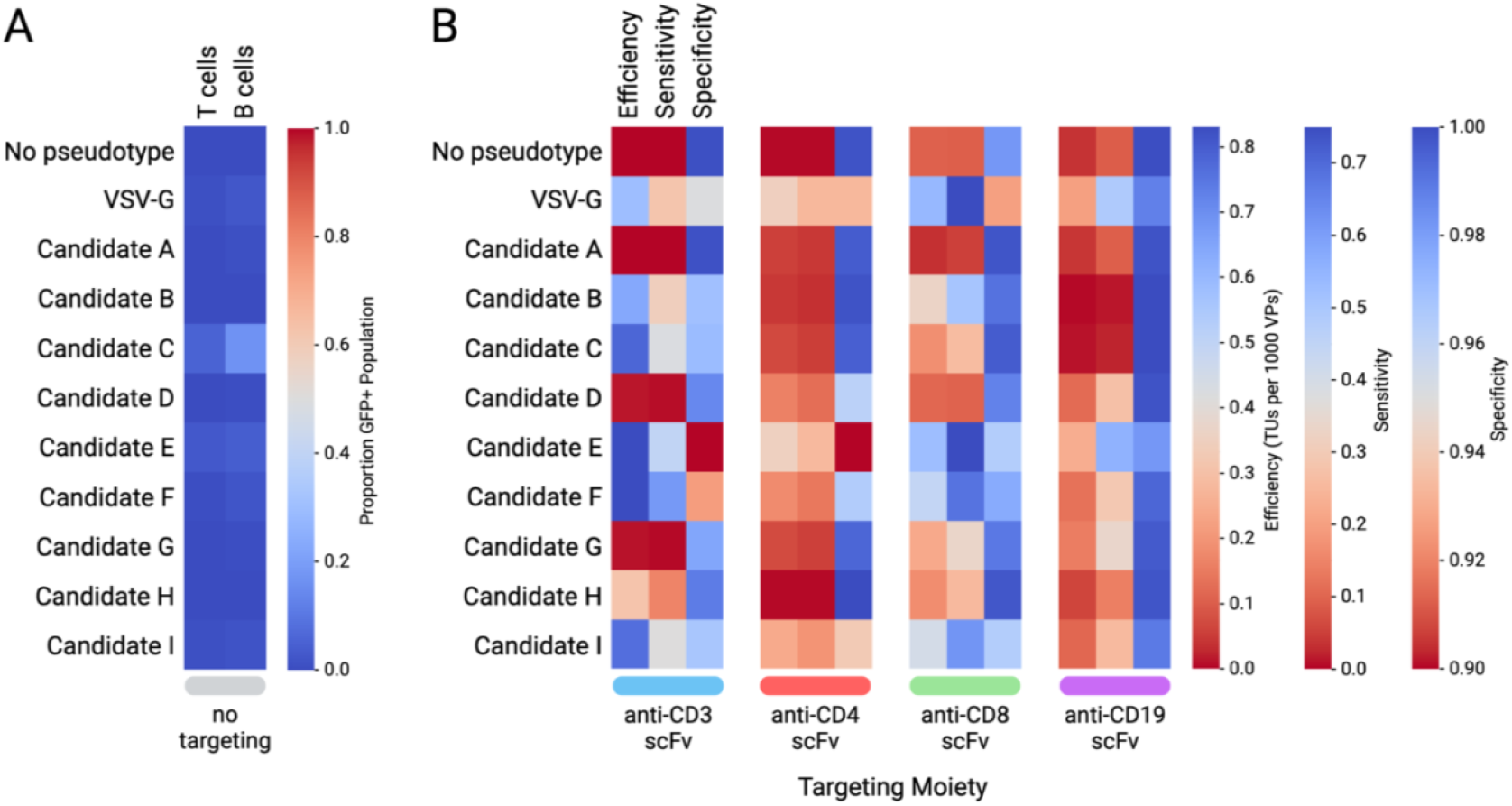
Medium-throughput screen of nine viral G-protein candidates for cell-targeted transduction of human PBMCs. (A) Proportion (%) of GFP positive T and B cells following transduction of human PBMCs with non-targeted lentivectors pseudotyped with the indicated viral G-proteins. (B) Transduction efficiency (TUs per 1000 VPs), sensitivity (%), and specificity (%) for lentivectors pseudotyped with the indicated viral G-proteins and grouped by the cell targeting moiety present on the lentivector.

To compare across the candidate viral G-protein and cell targeting combinations, we calculated efficiency of transduction (“efficiency”), sensitivity, and specificity values from the flow cytometry data (**Figure 3B; Supplementary Table S4**). To compare across pseudotypes and cell targeting moieties, we calculated on-target transduction efficiency (TUs/LP) using the PBMC transduction flow cytometry data to estimate the number of transduction units (TUs) for each lentivector sample and the associated lentivector particle (LP) count. Sensitivity is related to efficiency but is expressed as a percentage of on-target cells that receive the transgene rather than TUs/VP. Specificity is expressed as a percentage of off-target cells that do not receive the transgene. There is often a tradeoff between efficiency, sensitivity, and specificity which requires all these values to be considered when choosing the best option to move forward. For example, Candidate A had the highest specificity score (99.8%) but the lowest efficiency (3.4×10^−5^ TUs/LP) and sensitivity (4.7%) scores averaged across all scFvs tested with very few transduced target cells seen by flow cytometry. Conversely, VSV-G, Candidate E, and Candidate F had the highest efficiency (4.4×10^−4^, 5.3×10^−4^, and 5.1×10^−4^ TUs/LP, respectively) and sensitivity (46.3%, 51.6%, and 42.9%, respectively) scores, but the lowest specificity (95.0%, 91.0%, and 96.5%) scores averaged across all scFvs tested for each G-protein. The high efficiency and sensitivity, but lower specificity, scores for conventional VSV-G agrees with its longstanding use for efficient *in vitro* gene delivery to a diverse set of mammalian cell types and provides internal validation of our screen.

Across all lentivectors tested, efficiency was highest for the CD3 and CD8-targeted lentivectors (average 5.3×10^−4^ and 3.2×10^−4^ TUs/LP respectively). Similarly, CD3 and CD8-targeted lentivectors had the highest sensitivity scores (average 25.7% and 44.4%, respectively). These CD3 and CD8-targeted lentivector samples also had higher GFP signal strength than CD4 and CD19-targeted lentivector samples possibly due to more efficient cell entry and subsequent gene delivery to the target cells. Interestingly, some pseudotypes performed better when paired with certain cell targeting moieties. For example, Candidate B performed well for CD8-targeted gene delivery with 50.5% sensitivity and 99.1% specificity but poorly for CD19-targeted gene delivery with 1.28% sensitivity, whereas Candidate G performed well for CD19-targeted gene delivery with 33.8% sensitivity and 99.7% specificity but poorly for CD3-targeted gene delivery with 0.5% sensitivity.

For CD3-mediated gene delivery, multiple G-proteins mediated strong T cell transduction with variable levels of off-target transduction. Thus, we decided to move CD3-mediated lentivector designs into *in vivo* tests. Candidate F had the highest sensitivity (61.0%) with specificity of 92.6% followed by Candidate E with sensitivity of 44.8% and a low specificity of 80.8%. Candidates C (sensitivity: 38.1%: specificity: 97.1%) and I (sensitivity: 37.3%; specificity: 96.7%) had slightly lower sensitivity but higher specificity scores. Finally, Candidate B (sensitivity: 31.2%; specificity: 96.9%) and VSV-G (sensitivity: 28.4%; specificity: 95.0%) had reasonable sensitivity and specificity scores for CD3-mediated transgene delivery.

### In vivo *transduction efficiency pilot study in mice shows that three candidate viral G-proteins perform at least as well as conventional VSV-G*

We were interested to determine how the top *in vitro* performing viral G-protein candidates performed *in vivo* for cell targeted gene delivery to human T cells in a PBMC humanized immunodeficient mouse model (**Figure 4A**). For these *in vivo* pilot experiments, we conducted suspension packaging of lentivectors (**Figure 4B**) with a GFP transgene (**Figure 4C**), the anti-human CD3 scFv, and pseudotyped with VSV, mut-VSV, Candidate B, C, E, F, or I viral G-proteins, which comprise a range of sequence identities to VSV-G (40%, 41%, 38%, 38%, and 85%, respectively) and a range of CD3-targeted transduction sensitivities (31.2%, 38.1%, 44.8%, 61%, and 37.3%, respectively). The suspension-packaged lentivectors pseudotyped with Candidates E and F had low functional titers and were not moved forward for *in vivo* testing (**Supplementary Table S3**).

**Figure 4.**
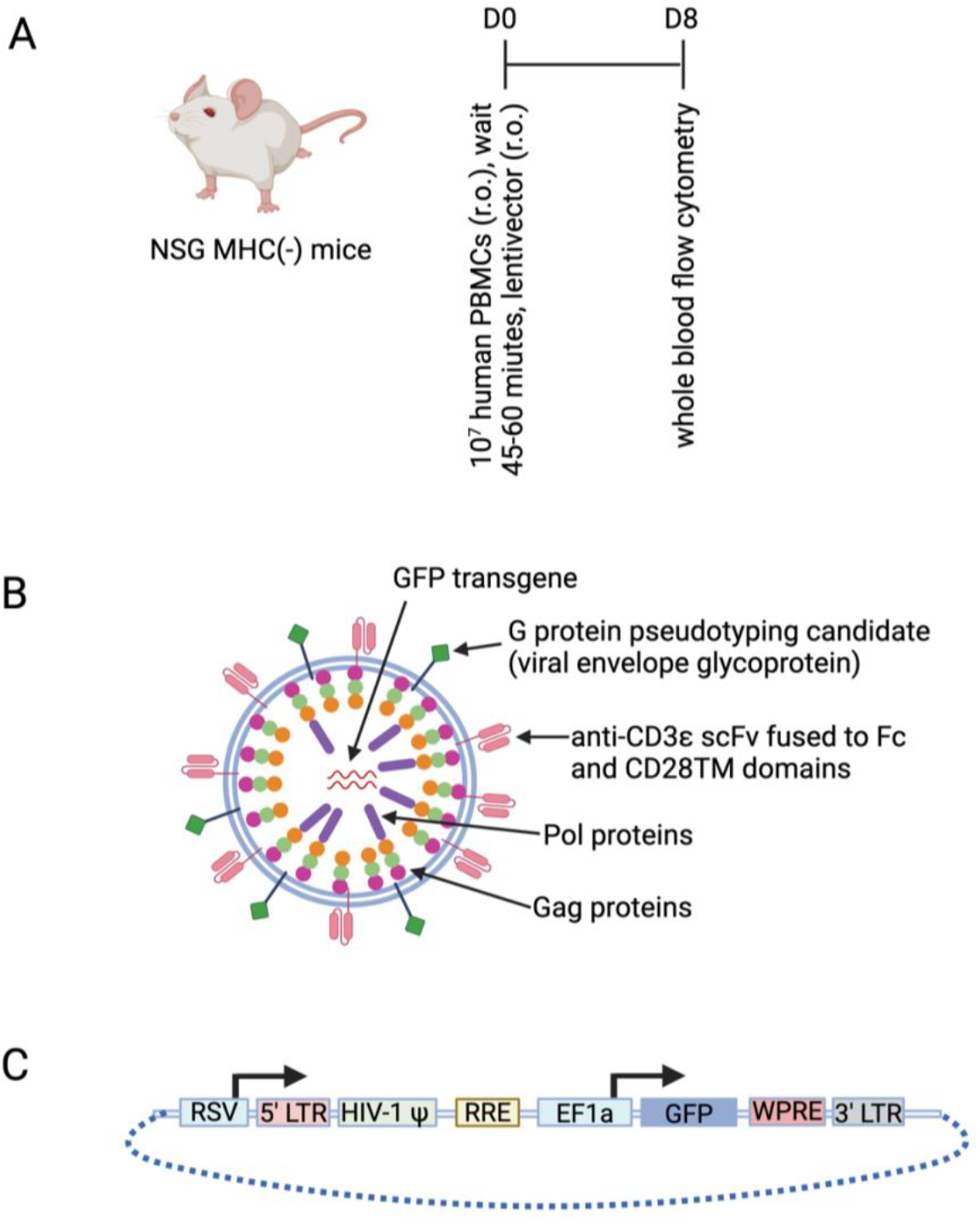
Pilot experiment to assess three candidate viral G-proteins for *in vivo* T cell-targeted gene delivery using human PBMC engrafted immunodeficient, NSG MHC DKO, mice. (A) Timeline for the pilot GFP transgene lentivector NSG MHC DKO mouse experiment, assessing three G proteins (Candidates B, C, and I), VSV-G, and mut-VSV-G, all directed to transduce T cells with an anti-human CD3 scFv. On Day 0 (D0), NSG MHC DKO mice were injected retro-orbitally (r.o.) into the right eye with 10^7^ human PBMCs. After 45-60 minutes, mice were injected r.o. into the left eye with one of the five lentivectors. Whole blood flow cytometry was performed on Day 8 (D8) to assess transduction efficiency. (B) General structure of the lentivectors used in the pilot GFP transgene mouse experiment. (C) General structure of the GFP transgene constructs used in the pilot GFP transgene mouse experiment. Elements in the transgene construct include: an RSV promoter (RSV), 5’ long terminal repeat (LTR), human immunodeficiency virus 1 psi packaging element (HIV-1 Ψ), rev response element (RRE), elongation factor 1 alpha (EF1a) promoter, DasherGFP (GFP) coding region, woodchuck hepatitis virus post-transcriptional response element (WPRE), and 3’ LTR.

NSG MHC DKO mice, which have reduced graft-versus-host disease when humanized, were injected r.o. with 10^7^ human PBMCs. 45-60 minutes later, mice were injected r.o. with 2×10^10^ LPs for one of the five lentivectors or DPBS vehicle control (n=3 mice per group). One week after injection, the percentage of circulating human T cells positive for GFP expression was measured (**Figure 5** and **Supplementary Figure S8**). GFP positive human T cells in mice injected with DPBS (vehicle) were undetectable, and there were no statistically significant differences in the percentage of GFP-positive human T cells among animals injected with lentivectors pseudotyped with Candidate B, Candidate C, or Candidate I (two-sample t-test, p>0.05). The only statistically significant difference between treatment groups was VSV-G vs. Candidate C (two-sample t-test, p=4.8×10^−4^), suggesting that the lentivector pseudotyped with Candidate C was more efficient at delivering transgenes *in vivo* than lentivectors pseudotyped with VSV-G. Thus, we proceeded with Candidate C for our subsequent *in vivo* tumor control study.

**Figure 5.**
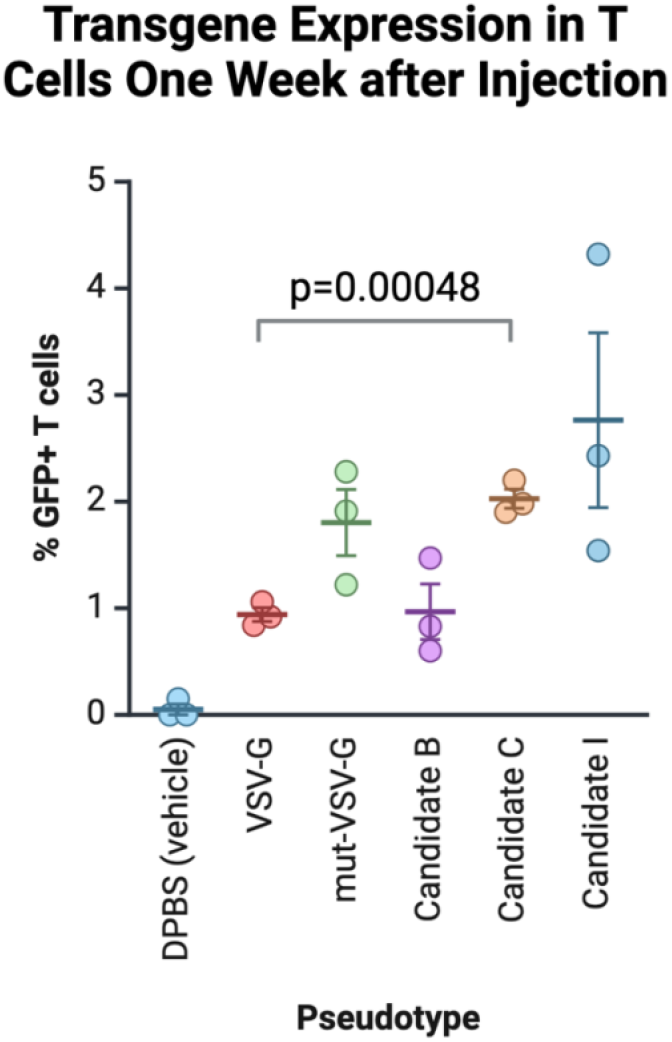
Transduction results from the pilot experiment to assess three candidate viral G-proteins for *in vivo* T cell-targeted gene delivery using human PBMC engrafted immunodeficient, NSG MHC DKO, mice. Shown are the percentages of human T cells positive for GFP expression in mice injected with DPBS (vehicle), and anti-human CD3 scFv directed lentivectors pseudotyped with VSV-G, mutated VSV-G (mut-VSV-G), Candidate B, Candidate C, or Candidate I. PBMCs were assessed by flow cytometry 8 days after injection. Error bar is standard deviation across replicate mice in each treatment group. P-value for VSV-G vs. Candidate C is a two-sample t-test. Error bar is standard deviation across replicates.

### Mouse NALM6 tumor model shows that a candidate viral G-protein can mediate efficient in vivo delivery of a CAR construct to human T cells resulting in tumor control

We performed a mouse tumor control study using immunodeficient, NSG MHC DKO, mice engrafted with ffLuc^+^ NALM6 B cell lymphoma cells by i.p. injection (**Figure 6A**). Mice were injected with luciferase-expressing NALM6 cells on day −14 (D-14), and whole body BLI was conducted on D-8 and D-1. Mice were randomized into treatment groups based on their BLI signal on D-1. On day 0 (D0) mice were injected r.o. with human PBMCs followed by lentivector 45-60 minutes later (**Figure 6B**). Group 1 animals (n=6) did not receive either PBMCs or lentivector. Group 2 animals (n=7) were injected with human PBMCs and human CD3-targeted Candidate C pseudotyped lentivector with a GFP transgene. Group 3 animals (n=7) were injected with human PBMCs and human CD3-targeted Candidate C pseudotyped lentivector with an anti-CD19 CAR-GFP transgene (**Figures 6C-D**). Mouse weights were taken twice weekly (**Supplementary Figure S9**) and whole body BLI (**Supplementary Figure S10**) and flow cytometry on mouse whole blood samples was conducted weekly over the course of the study. Human PBMCs grafted stably in Groups 2 and 3 over the course of the study (**Supplementary Figure S11**).

**Figure 6.**
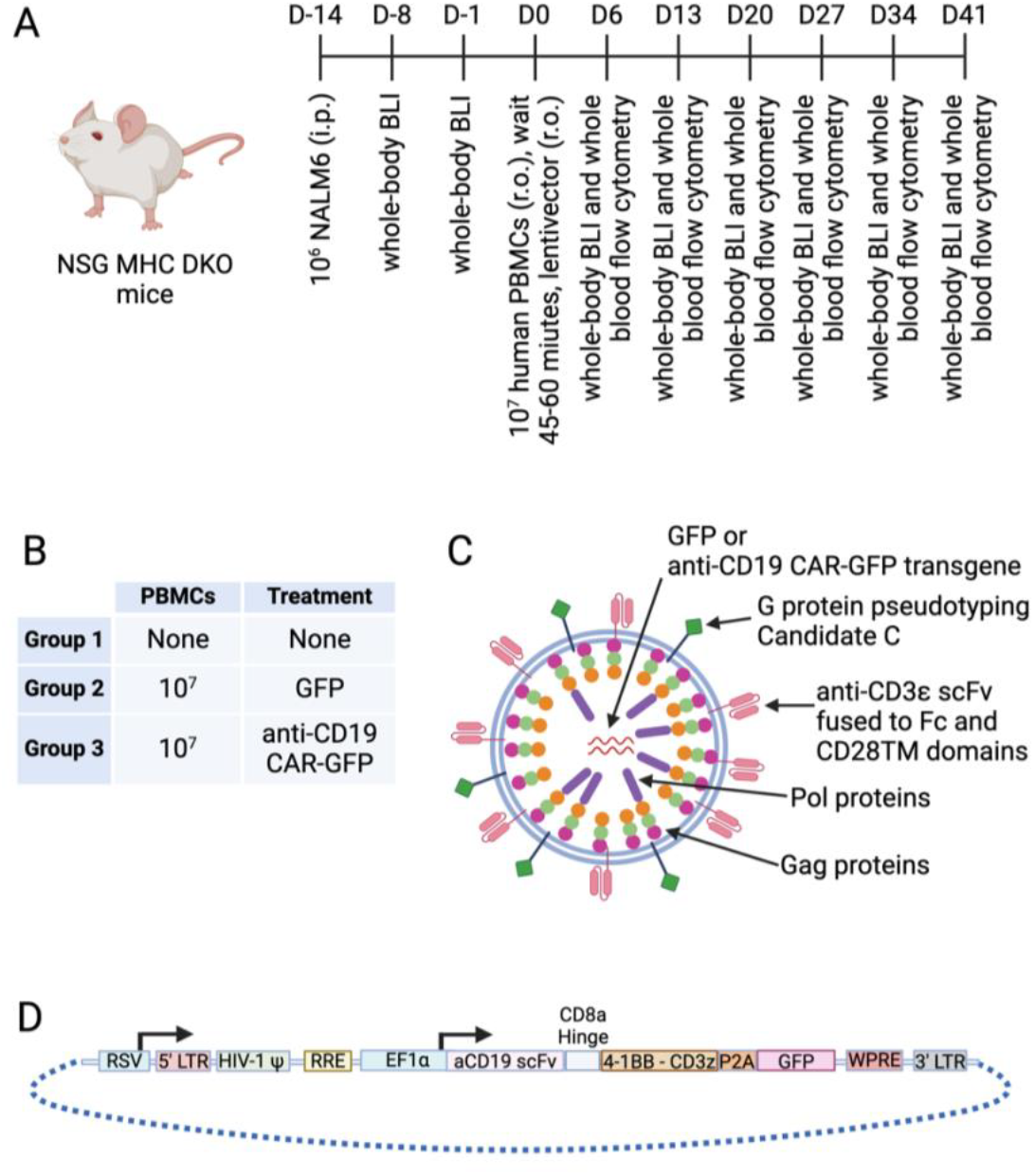
Tumor control study of a lentivector bearing an anti-CD19 CAR for in vivo T cell-targeted gene delivery using human PBMCs engrafted NSG MHC DKO immunodeficient mice. (A) Timeline for the NALM6 tumor control study in NSG MHC DKO mice, assessing viral G-protein Candidate C with an anti-human CD3 scFv. On Day −14 (D-14), mice were injected with 10^6^ firefly luciferase-engineered NALM6 cells intraperitoneally (i.p.). Whole-body bioluminescent imaging (BLI) was performed on Days −8 and −1 (D-8 and D-1), and mice were randomized by luminescence signals on D-1. On Day 0 (D0), mice were injected r.o. into the right eye with 10^7^ human PBMCs (Group 2, n=7; and Group 3, n=7). 45-60 minutes later, GFP transgene lentivector (Group 2), or anti-CD19 CAR-GFP transgene lentivector (Group 3), was injected r.o. into the left eye. Group 1 mice (n=6) did not receive PBMC or lentivector injections. Whole blood flow cytometry and whole-body BLI were performed weekly through the end of the study. (B) Treatment groups in the NALM6 tumor control study. (C) General structure of the lentivectors used in the mouse tumor control study. (D) General structure of the anti-CD19 CAR-GFP transgene constructs used in the mouse tumor control study. Elements in the transgene construct include: an RSV promoter (RSV), 5’ long terminal repeat (LTR), human immunodeficiency virus 1 psi packaging element (HIV-1 Ψ), rev response element (RRE), elongation factor 1 alpha (EF1a) promoter, a multicistronic anti-CD19 scFv – CD8a hinge – 4-1BB-CD3z CAR plus DasherGFP (GFP) coding region with a P2A ribosome skip motif, woodchuck hepatitis virus post-transcriptional response element (WPRE), and 3’ LTR.

Animals injected with GFP-only lentivector (Group 2) had an average of 2.5% GFP^+^ human T cells at 1 week (range: 1.23% to 3.29%) and was stable over the course of the study (**Figure 7A**). While the average percent of CAR-GFP^+^ human T cells in Group 3 was only 0.3% (range: 0.17% to 0.48%) at 1 week, this increased to an average of 4% (range: 0.9% to 9.0%) on week 2 and peaked at an average of 20% (range: 4.9% to 39.3%) on week 3 (**Figure 7A**). By Day 20, there was a statistically significant difference between the GFP signal in Group 2 and Group 3 (two-sample t-test, p=0.003). GFP signal was not detected in mouse CD45^+^ cells at any timepoint, and only a single mouse sample (MIV-020-0007) from Group 2 (GFP only) showed GFP^+^ cells at about 6% frequency in the human B cell population on D41, the last day of the study. Note this mouse had a massive tumor burden at this timepoint and its unclear if this population of GFP^+^ human B cells would have persisted to later timepoints. These data suggest highly specific *in vivo* delivery to CD3^+^ T cells.

**Figure 7.**
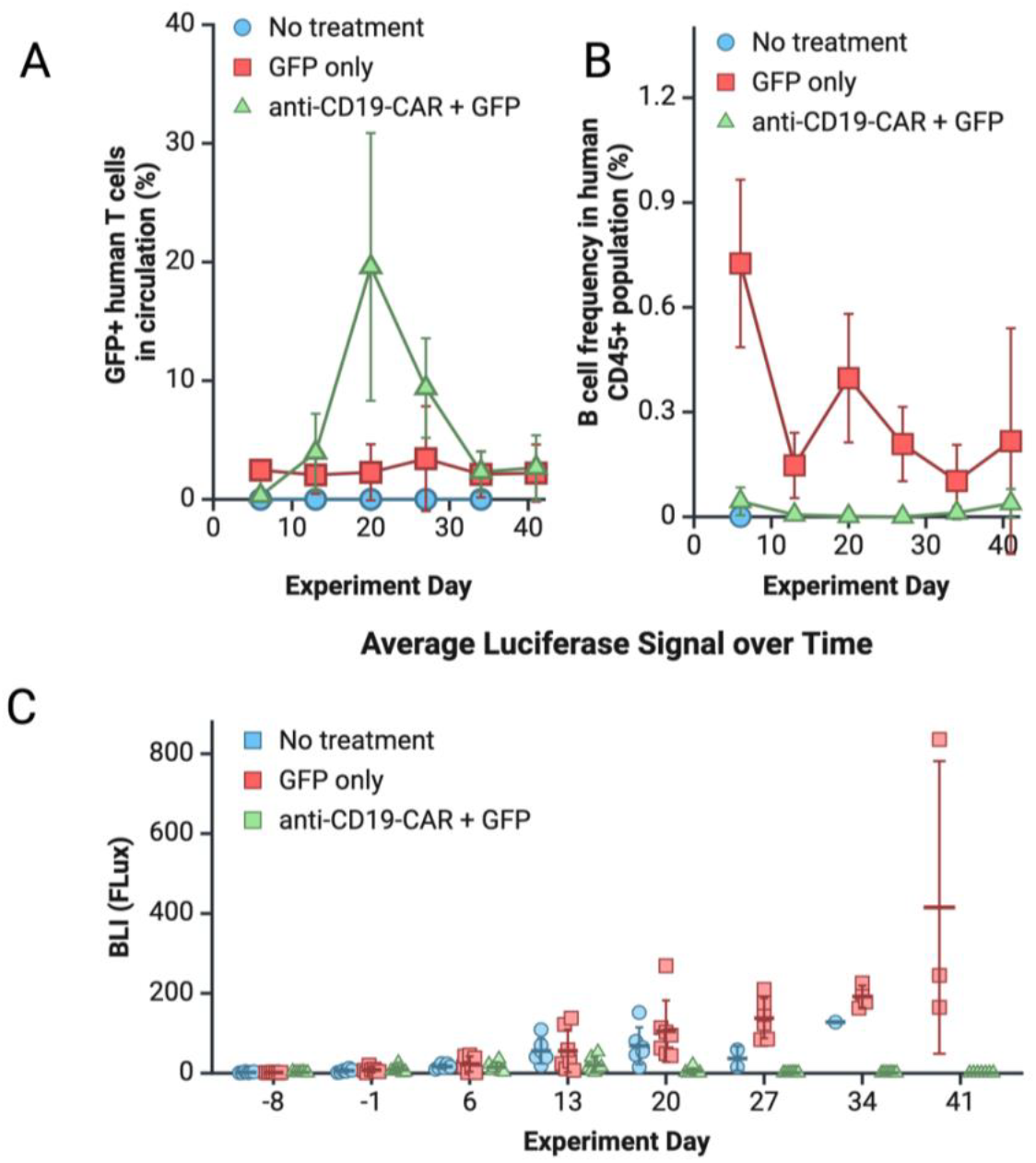
Pharmacology and tumor control following *in vivo* dosing of T cell-targeted lentivector bearing an anti-CD19 CAR-GFP transgene into NALM6-engrafted NSG MHC DKO mice humanized withPBMCs versus negative control arms. (A) Percentage of human T cells positive for GFP expression in mice injected with a GFP-only transgene (Group 2) or an anti-CD19 CAR-GFP transgene (Group 3), assessed by flow cytometry using whole blood samples on Days 6, 13, 20, 27, 34, and 41 after treatment with lentivector. Average values and standard deviation (error bars) across replicates are plotted. (B) Percentage of human B cells (CD19^+^,CD20^+^) in mice injected with a GFP-only transgene (Group 2) or an anti-CD19 CAR-GFP transgene (Group 3), assessed by flow cytometry using whole blood on Days 6, 13, 20, 27, 34, and 41 after treatment with lentivector. Average values and standard deviation (error bars) across replicates are plotted. (C) Average bioluminescence imaging (BLI) signal, reported as flux, across replicate mice in each treatment group, assessed one week (Day −8) after injection of NALM6 cells through eight weeks after injection of NALM6 cells (Day 41). Error bar is standard deviation across replicates.

*In vivo* delivery of the anti-CD19 CAR-GFP transgene to human T cells mediated a rapid depletion of PBMC derived B cells (**Figure 7B**). At week 1 post-treatment, Group 3 (anti-CD19 CAR-GFP transgene) mice had an average of 0.04% human B cells (range: 0.00% to 0.11%) compared to 0.73% frequency (range: 0.41% to 1.12%) in Group 2 (GFP transgene) mice, a statistically significant difference (two-sample t-test, p=0.0001). This suggests rapid killing of human B cells by anti-CD19 CAR-T cells *in vivo*.

Group 3 (anti-CD19 CAR-GFP transgene) mice demonstrated complete tumor control in the NALM6 model. Group 3 animals were healthy without signs of toxicity throughout the study and had no sign of tumor cells at the end of the study (D41), whereas Group 1 and 2 animals were either dead or had massive tumor burdens at the end of the study (**Figures 8-9**). In Groups 1 and 2, the average luciferase signal (i.e., tumor load) increased by 11.8-fold and 17.1-fold, respectively, between Day −1 and Day 27 (**Figure 7C**), indicating lack of tumor control. In contrast, tumor growth was controlled in Group 3 mice with no increase in luciferase signal observed between Days 6 and 13, a rapid decline in luciferase signal by Day 27 (an average of 90% decrease in signal between Days −1 and 27), and no detectable luciferase signal in any mouse by Day 27 (**Figure 8** and **Supplementary Figure S10**).

**Figure 8.**
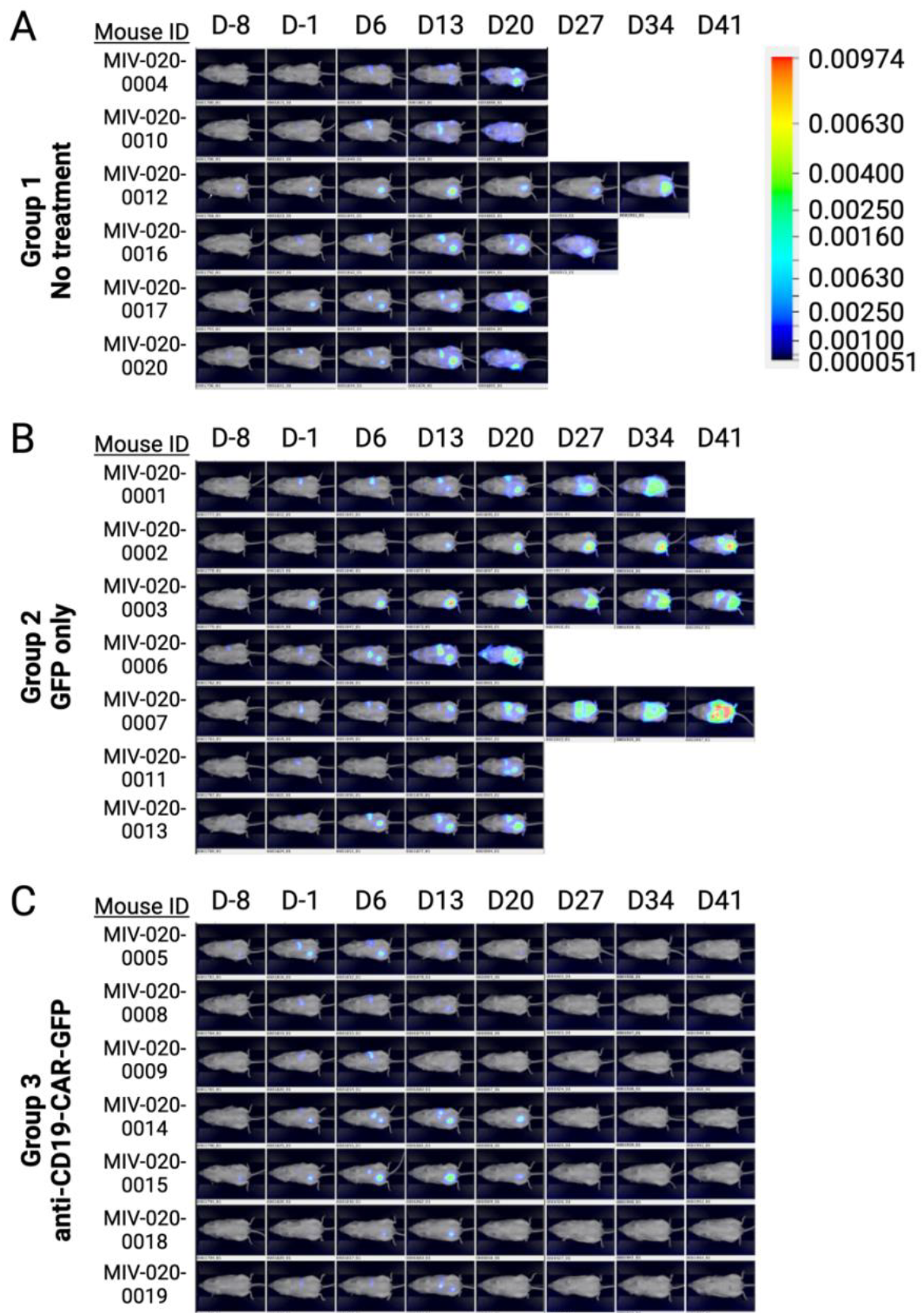
Tumor control following *in vivo* dosing of T cell-targeted lentivector bearing an anti-CD19 CAR-GFP transgene into NALM6-engrafted NSG MHC DKO mice humanized withPBMCs versus negative control arms. Luciferase signal detected by BLI in (A) Group 1 mice receiving no treatment, neither human PBMCs nor lentivector, (B) Group 2 mice injected with human PBMCs and GFP-only transgene, and (C) Group 3 mice injected with human PBMCs and anti-CD19 CAR-GFP transgene. Mice were imaged one week (D-8) through eight weeks (D41) after injection of NALM6 cells. Human PBMCs and lentivector dosing was completed on D0. Note, mouse MIV-020-0002 was euthanized on D41 following BLI due to poor body condition.

All Group 1 animals were dead by Day 41 (**Figure 9**) whereas two Group 2 animals remained alive at Day 41, both with massive tumor burden (**Figure 8**). No animals in Group 3 died. There was a significant difference (log rank test, p=0.0002) in survival between Group 1 (no treatment) and Group 3 (anti-CD19 CAR-GFP). There was also a significant difference (log rank test, p=0.007) between Group 2 (GFP transgene) and Group 3 (anti-CD19 CAR-GFP). We conclude that a single injection of CD3-targeted Candidate C pseudotyped lentivector delivering an anti-CD19 CAR-GFP transgene can effectively eliminate tumors in a NALM6 B cell lymphoma mouse model.

**Figure 9.**
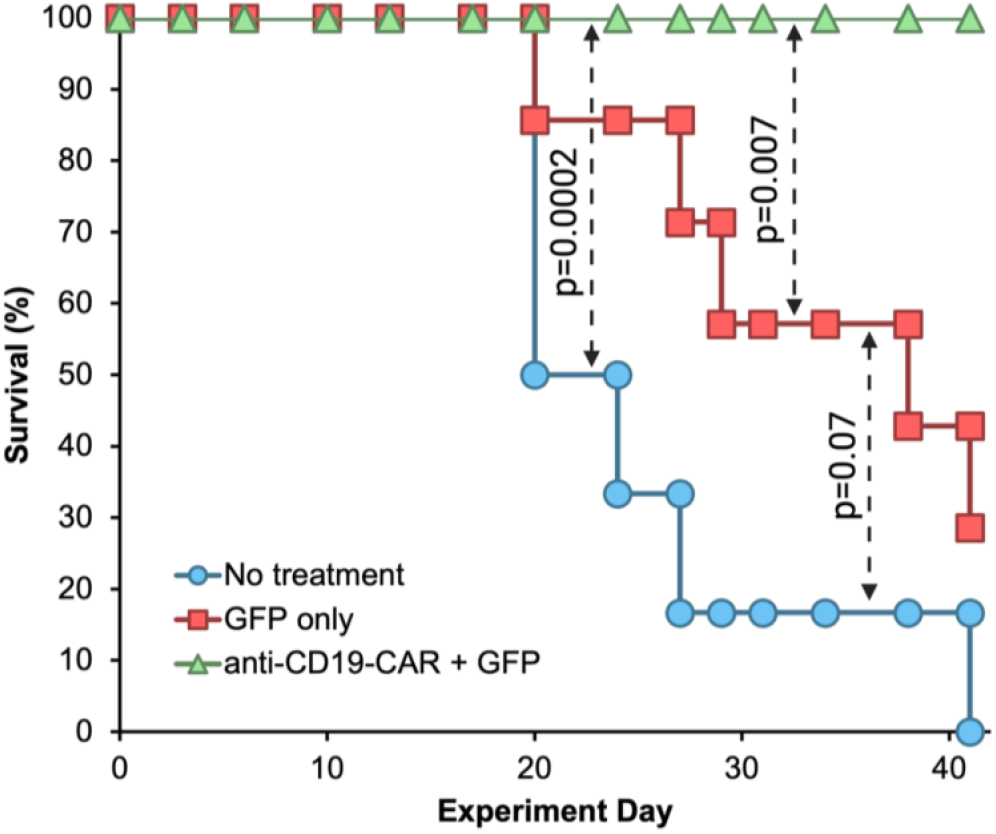
Survival curves for the NALM6 tumor control study treatment groups: no treatment (blue), human PBMCs and a single dose of lentivector carrying a GFP transgene (red), human PBMCs and a single dose of lentivector carrying an anti-CD19 CAR-GFP transgene (green). Indicated p-values were calculated using a log rank test.

## DISCUSSION

We see many opportunities for future research and optimization in our *in vitro* methods. Data generated with high-throughput screening methods like our *in vitro* screen in PBMCs are often less accurate than data generated using methods that are less high-throughput. We could have improved data quality by making replicate lentivector preparations. Pseudotransduction (Liu et al., 1996), a phenomenon where transgenes are transiently expressed without target genome integration, may have artificially increased the efficiency measurements in the *in vitro* screen in PBMCs. Nucleic acid barcoding and deep sequencing (e.g., Illumina) could be used to screen for transduction efficiency of large lentivector libraries *in vitro*. Such methods would allow for testing of transduction using millions of combinations, for example, diverse libraries of scFvs and/or pseudotypes. Such methods could be used to screen for features other than efficiency, specificity, and sensitivity, for example, heat stability (by incubating libraries at high temperatures prior to transduction) or immunogenicity (by incubating libraries in serum prior to transduction).

Our *in vivo* models were complicated, but commonly used in the field (e.g., Michels et al., 2023) and designed to be translatable to human studies. Using fully human PMBCs and a human tumor model ensured that the molecular targets were relevant to human studies. However, our models do have some downsides, for example, allogeneic tumor killing by grafted human PBMCs, and natural decrease in B cell counts over time. Syngeneic tumor models can be designed in a way that allogeneic reactivity does not happen, and B cell populations are typically stable in wild type animals. Certainly, our methods could be repeated using other B cell lymphoma models, which may, for example, be more aggressive or suffer from antigen escape. We could also test our methods with other tumor targets, for example CD20 rather than CD19, or we could test our methods with other T cell targeting moieties on the lentivector, such as CD7 or CD8. However, we achieved our primary goal of showing proof-of-concept in a widely accepted translational cancer model.

Our ultimate goal is to bring *in vivo* lentivectors into clinical practice for the treatment of cancer. At the time of writing, we are not aware of any published data showing intravenous clinical administration of lentivectors. Still, our approach could have important advantages for the field. First, alternative pseudotypes should allow for subsequent doses of the same, or different, CAR transgenes. For example, if a drug developer delivers anti-CD19 CAR with a lentivector pseudotyped with VSV-G, but the patient relapses, a subsequent dose of lentivector pseudotyped with Candidate C could be used to deliver an anti-CD20 CAR transgene. Second, further development efforts may find that VSV-G, or similar G-proteins, are not sufficiently specific to T cells or manufacture poorly. A larger portfolio of pseudotypes could help solve such problems. We are optimistic that *in vivo* delivery of lentivectors is a promising approach for the future, and that optimized search, discovery, and development methods such as ours will be crucial to establish the approach in routine clinical practice.

## ACKNOWLEDGMENTS

Thanks to Drs. Jan Fredrik Simons, Yoong Wearn Lim, and Mark Robbins for helpful comments on study design, implementation, and interpretation. The staff at Mispro performed some cheek blood draws, daily cage checks, IACUC review, and veterinary care for the mouse studies. Bettina Iliopoulou provided technical support for mouse studies. All figures were prepared with Biorender.

## SUPPLEMENTARY FIGURES

**Supplementary Figure S1.**
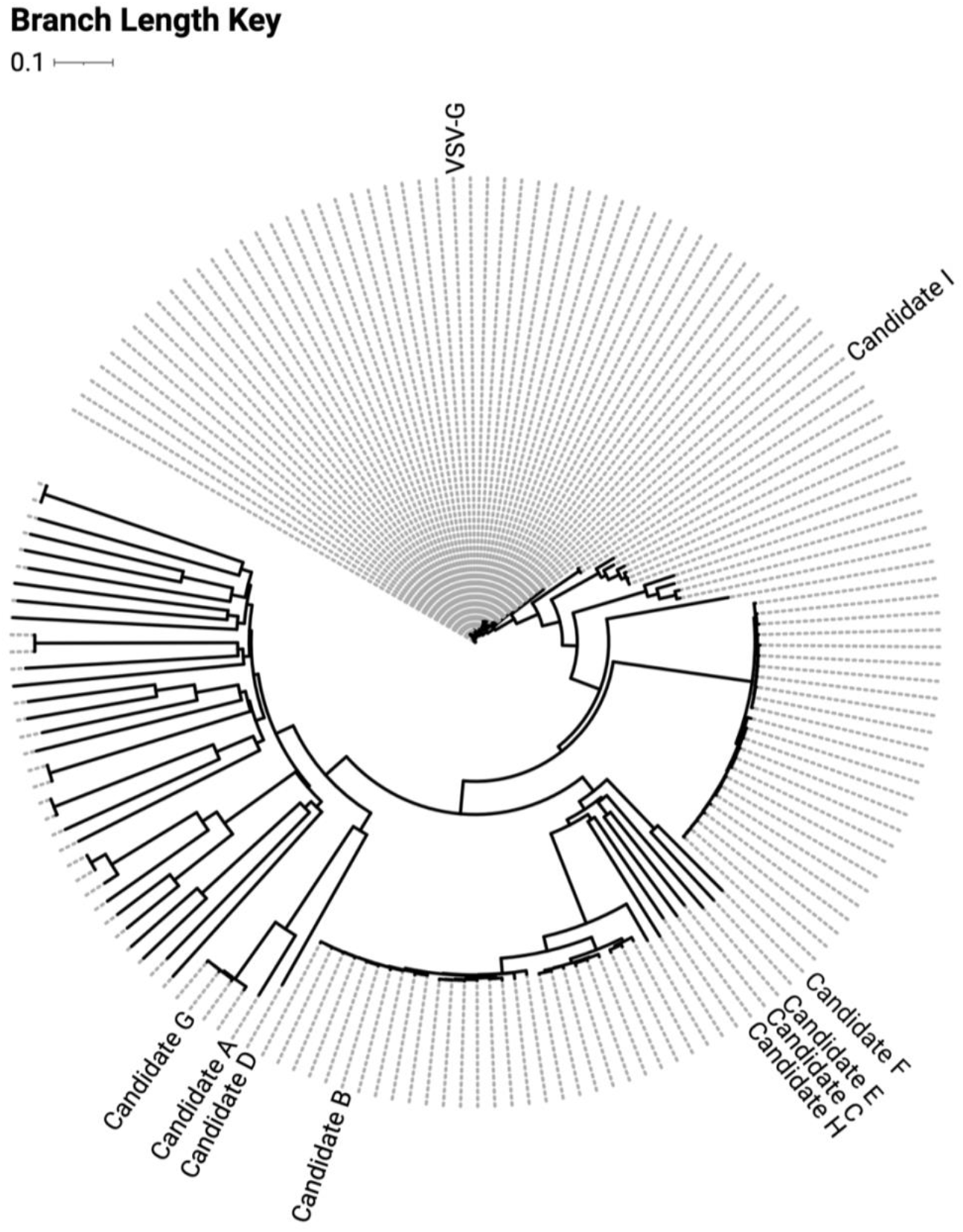
Phylogenetic tree for the 166 VSV-G-like G proteins identified in the Serratus search. The phylogenetic tree was generated with CLUSTAL-O and rendered with iTOL version 7.0 (Letunic & Bork, 2024). Branch lengths are indicated with solid black lines. Tree is rooted using the midpoint method. Nodes corresponding to VSV-G and Candidates A through I are labeled accordingly.

**Supplementary Figure S2.**
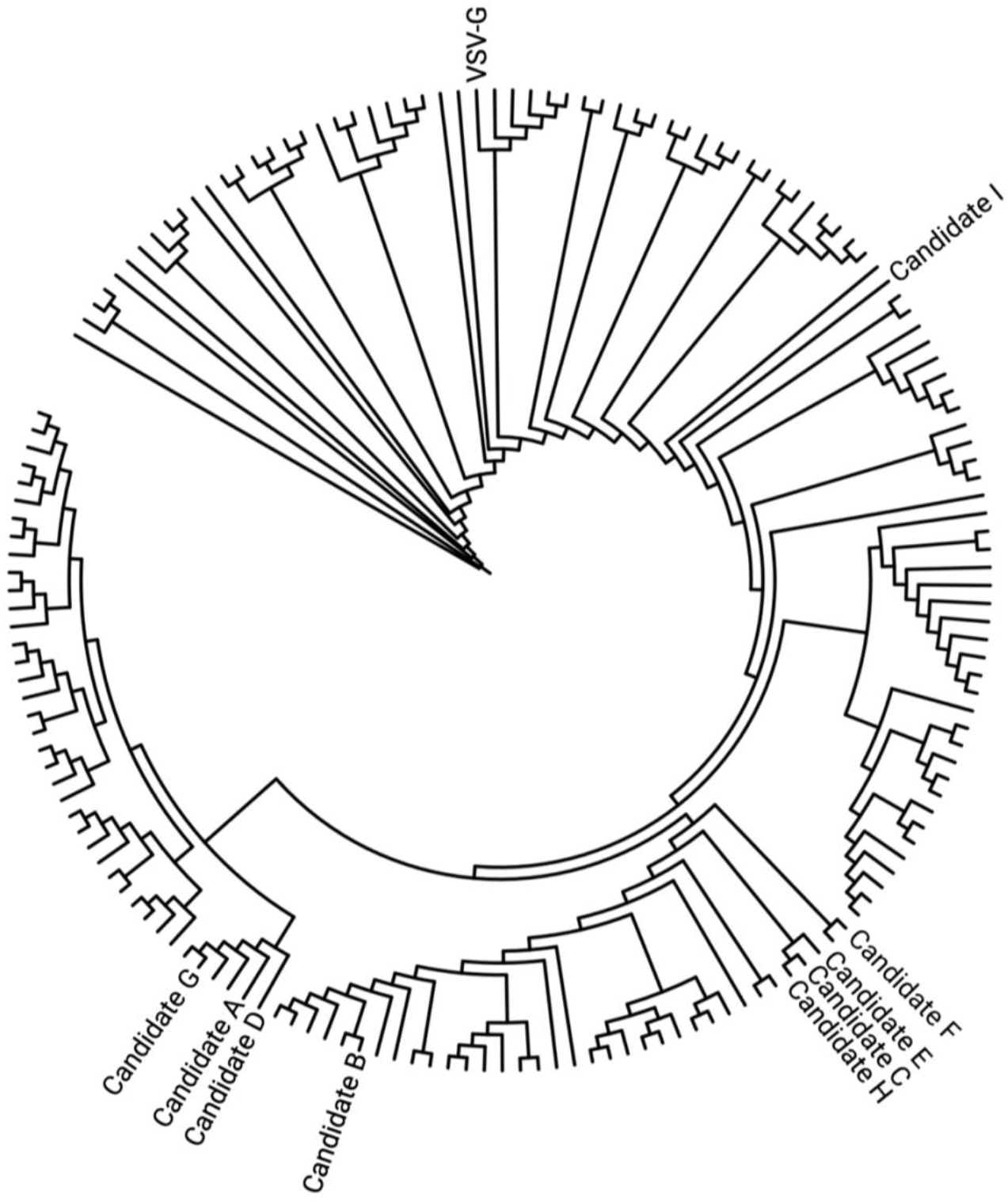
Phylogenetic tree for the 166 VSV-G-like G proteins identified in the Serratus search. The phylogenetic tree was generated with CLUSTAL-O and rendered with iTOL version 7.0 (Letunic & Bork, 2024). Tree is rooted using the midpoint method. Branch lengths are ignored. Nodes corresponding to VSV-G and Candidates A through I are labeled accordingly.

**Supplementary Figure S3.**
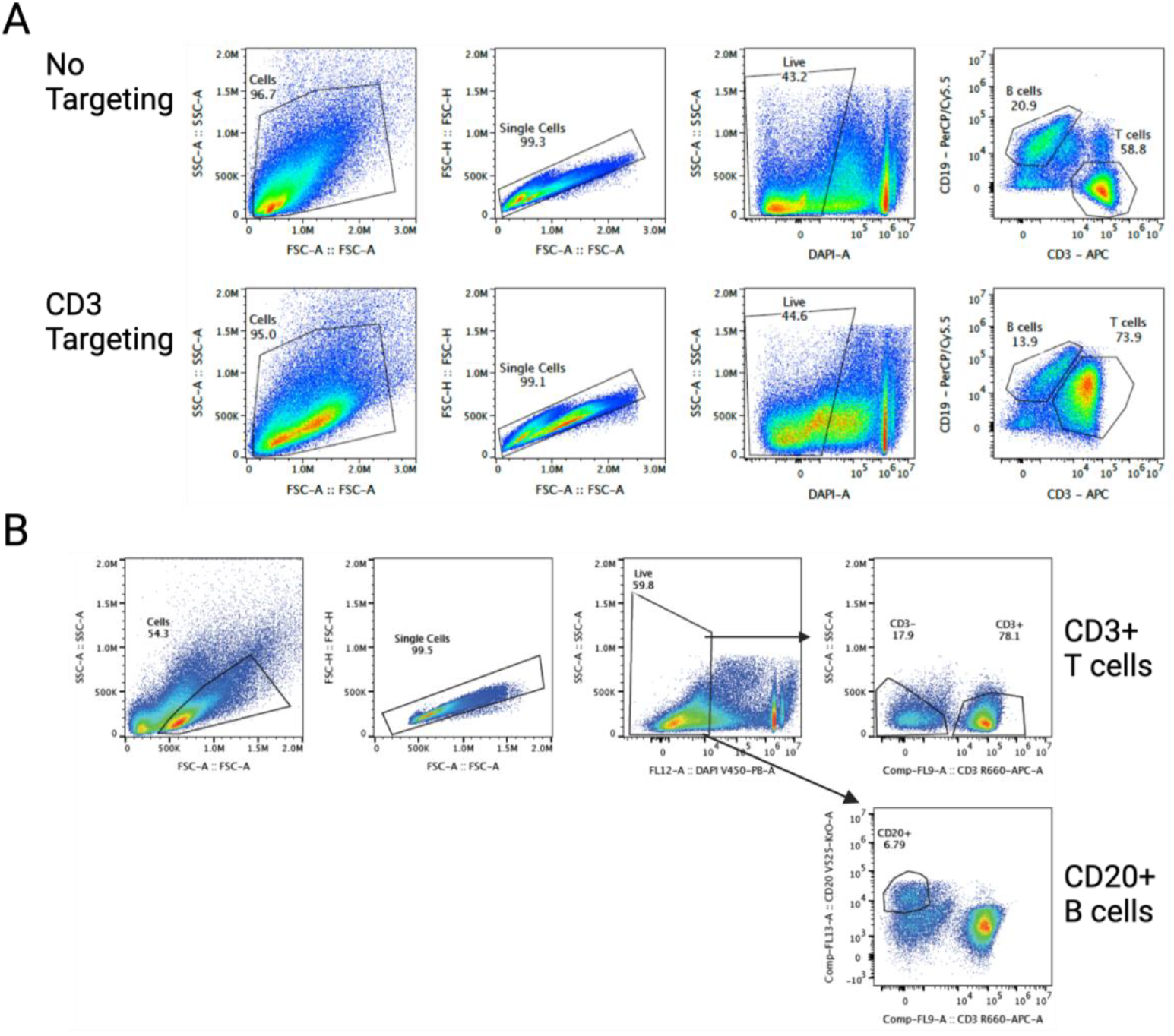
Flow cytometry gating strategies for pilot cell-targeted lentivector transduction of human PBMCs. (A) PBMC transduction with non-targeted and CD3-targeted lentivectors pseudotyped with VSV-G, mut-VSV-G, Candidate A, B, or C. Associated with **Figure 2A**. (B) PBMC samples transduced with Candidate C pseudotyped lentivector with anti-CD19, anti-CD4, or anti-CD8 scFv targeting. Associated with **Figure 2B**.

**Supplementary Figure S4.**
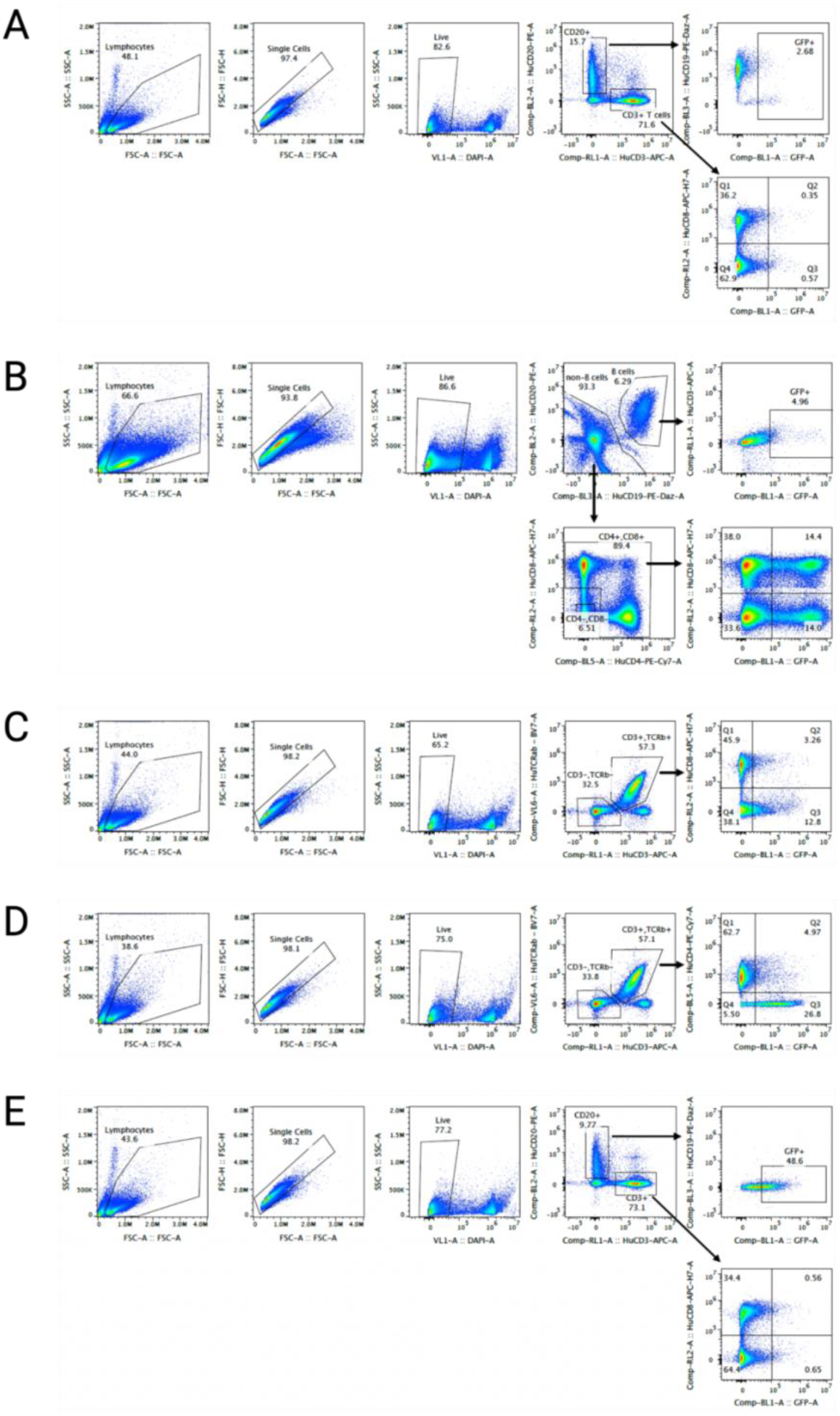
Flow cytometry gating strategies for the pseudotype by cell targeting moiety combination screen in human PBMCs. (A) No targeting. (B) CD3 targeting. (C) CD4 targeting. (D) CD8 targeting. (E) CD19 targeting. Associated with **Figure 3**.

**Supplementary Figure S5.**
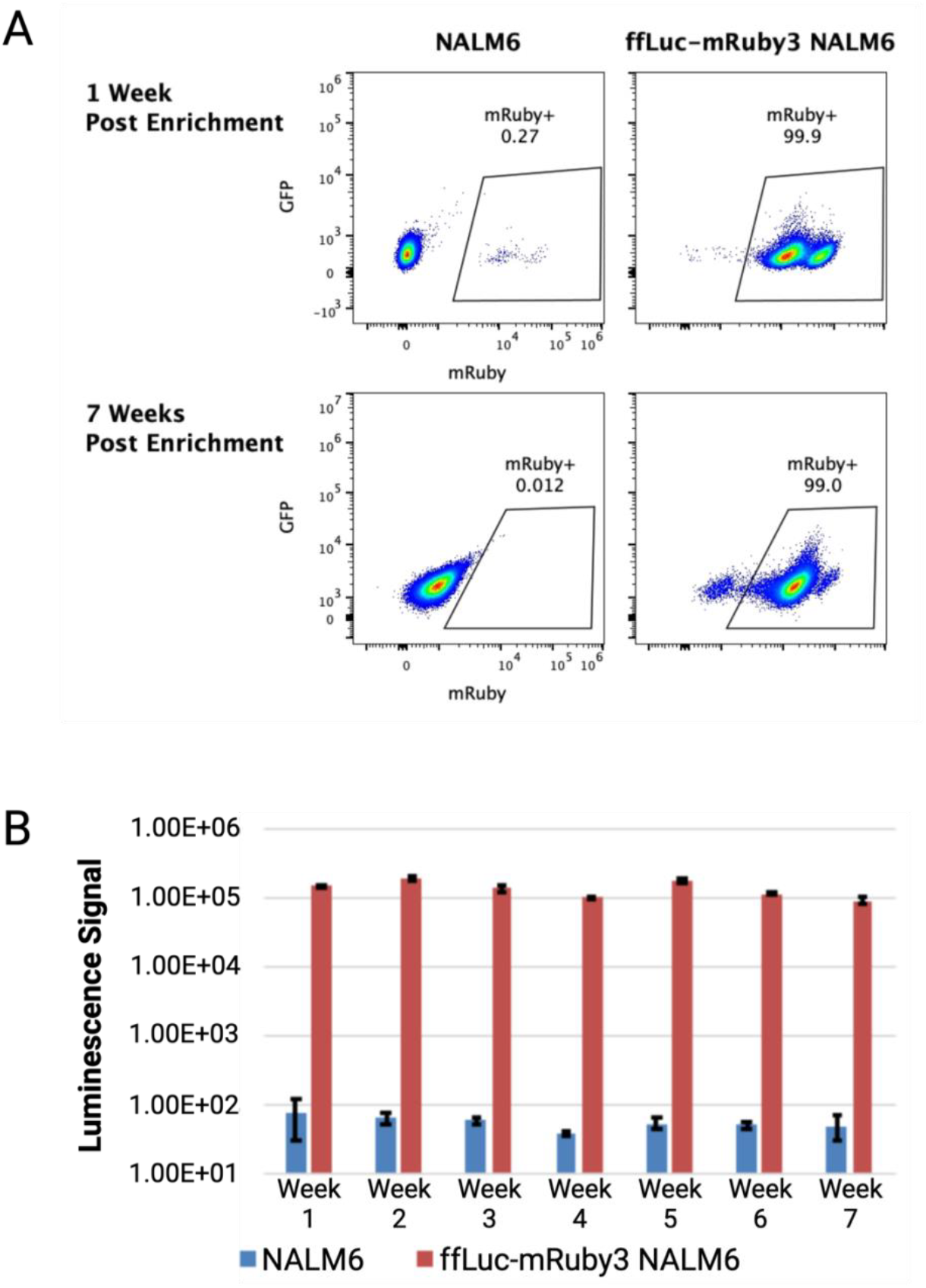
Firefly luciferase (ffLuc)-mRuby3 NALM6 cell line characterization. (A) Flow cytometry of parental and ffLuc-mRuby3 NALM6 cells 1 and 7 weeks after FACS enrichment for mRuby3+ cells. (B) Average bioluminescence signal from quadruplicate wells for parental and ffLuc-mRuby3 NALM6 cells over 7 weeks of culture. Error bars are standard deviation across replicates.

**Supplementary Figure S6.**
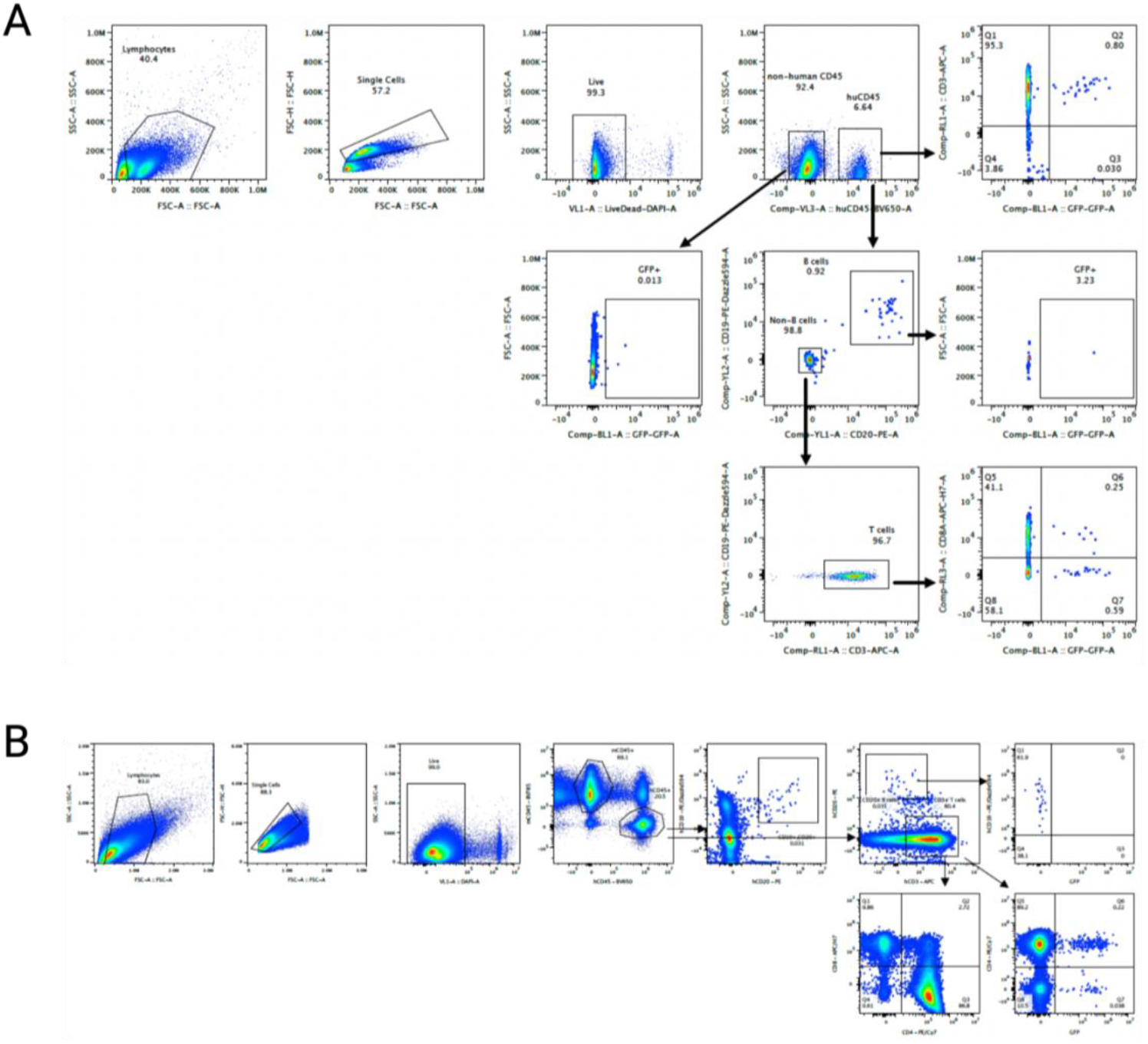
Flow cytometry gating strategies for (A) pilot *in vivo* transduction efficiency study, associated with **Figure 5**, and (B) mouse *in vivo* tumor control study, associated with **Figure 7**.

**Supplementary Figure S7.**
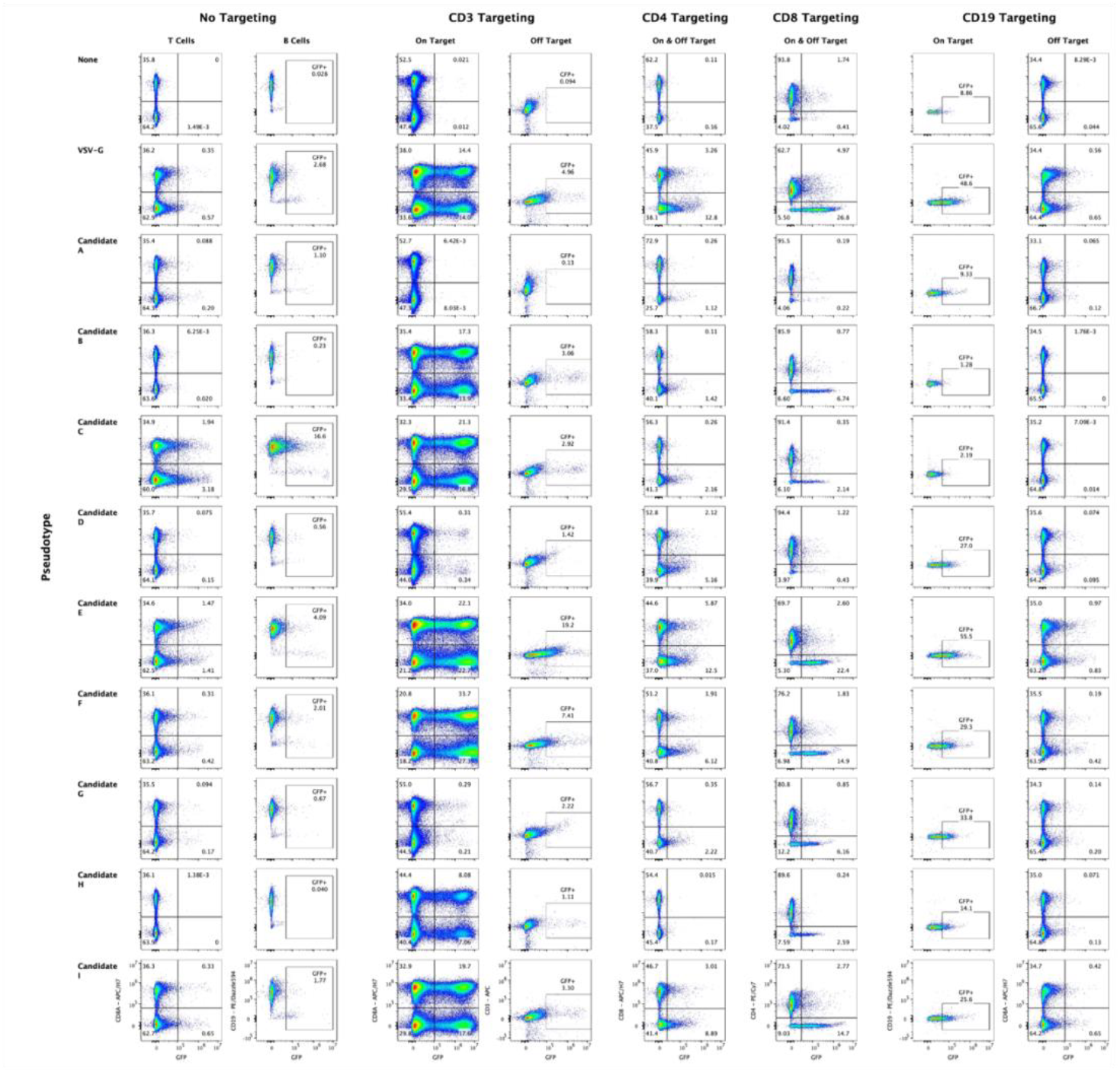
Flow cytometry data from the *in vitro* screen of nine viral G-protein candidates for cell-targeted transduction of human PBMCs. Associated with **Figure 3**.

**Supplementary Figure S8.**
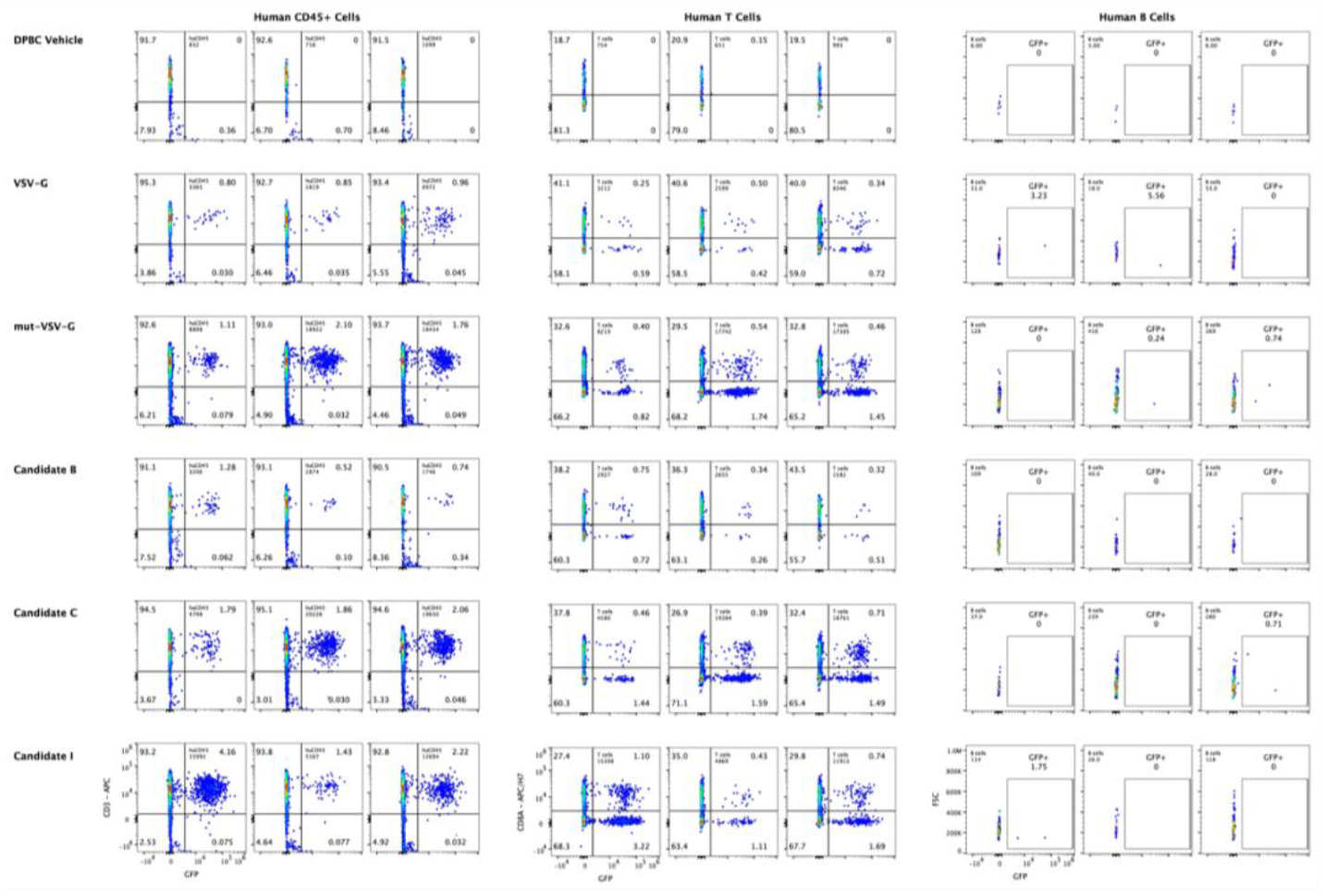
Flow cytometry data from the *in vivo* transduction efficiency pilot study. Associated with **Figure 5**.

**Supplementary Figure S9.**
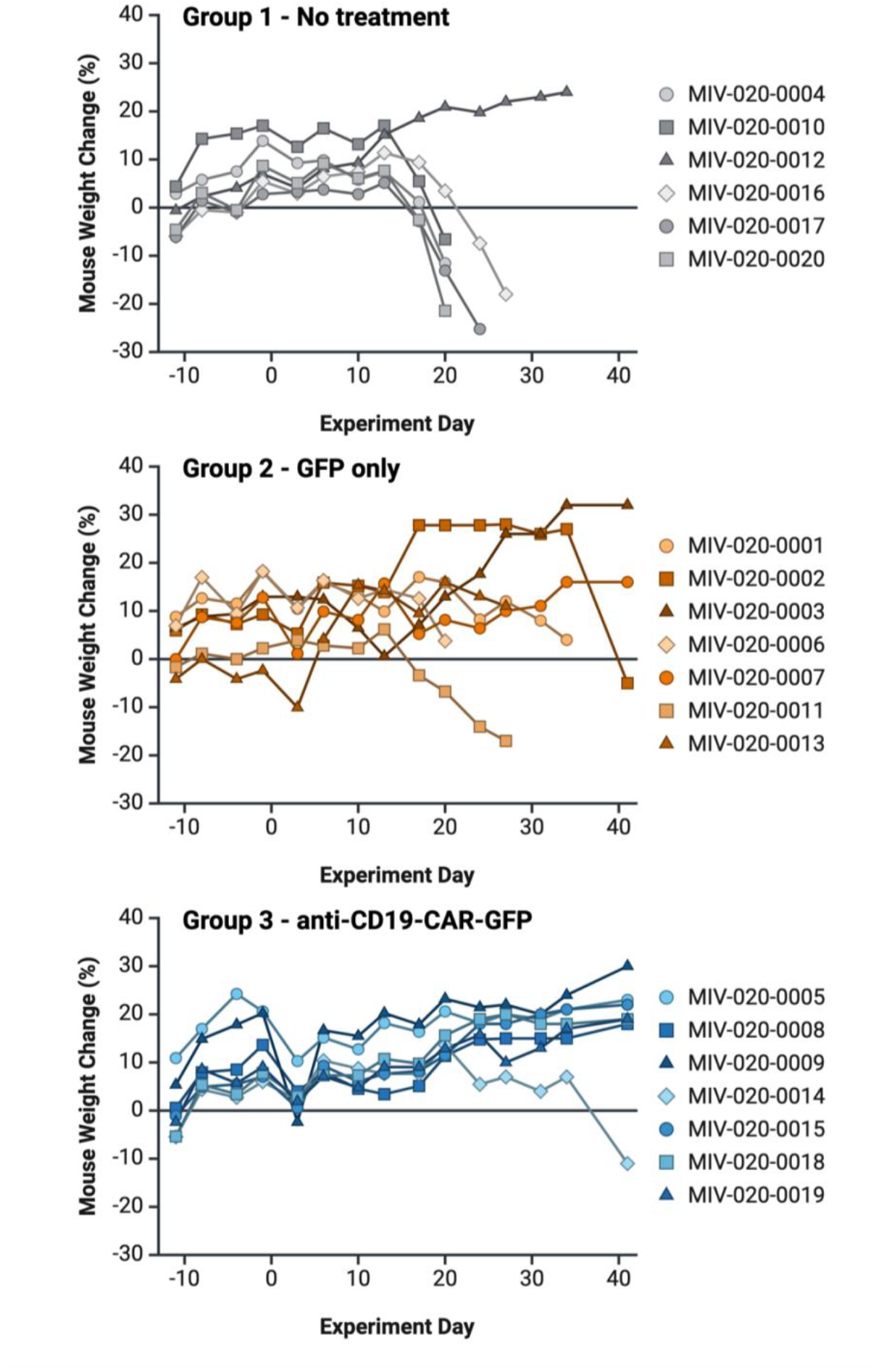
Mouse weights, with a sampling of days from Day −11 through Day 34, for individual mice in Groups 1-3 in the mouse *in vivo* tumor control study.

**Supplementary Figure S10.**
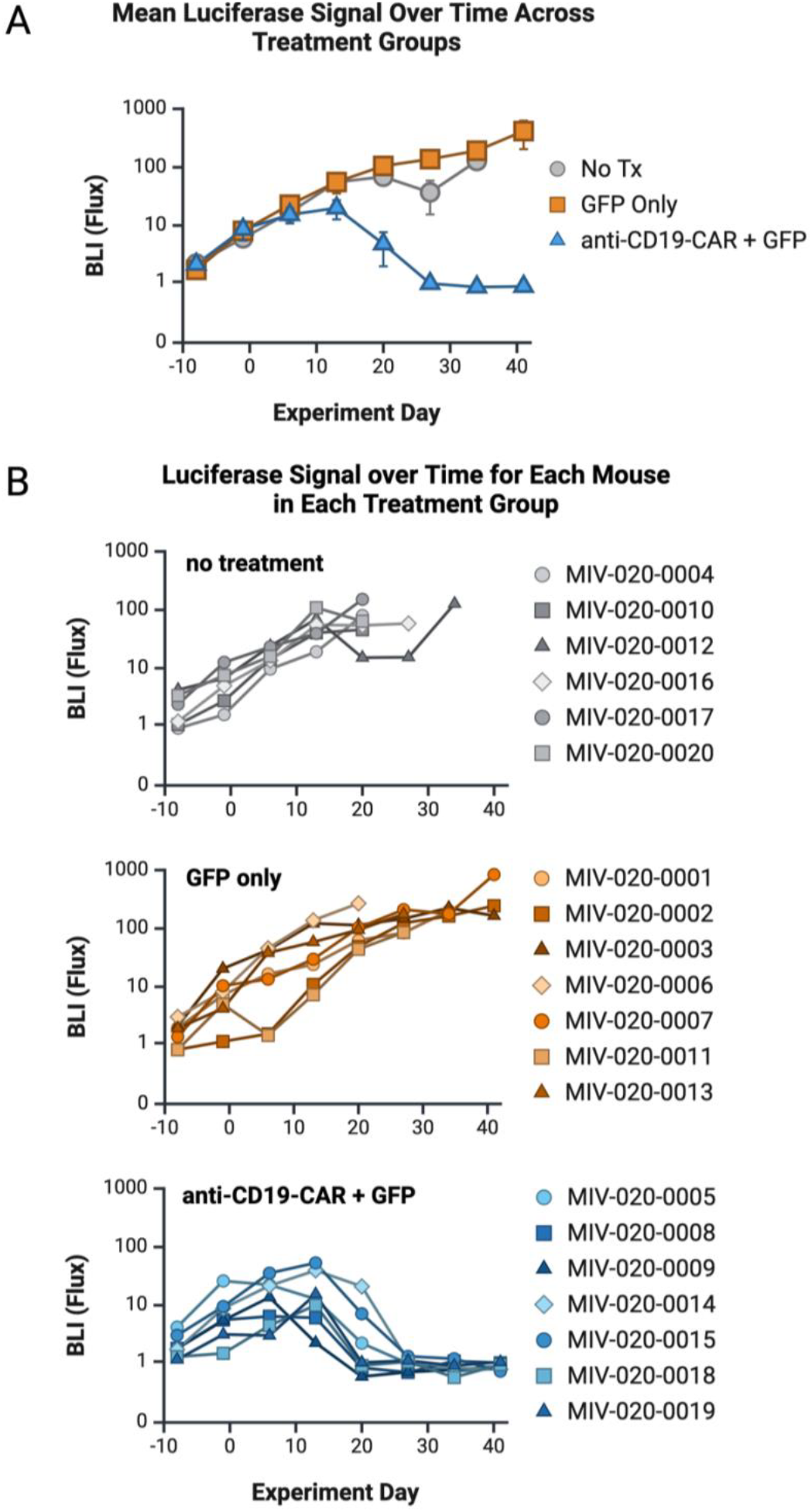
(A) Average BLI (flux) across mice withing each treatment group of the mouse *in vivo* tumor control study, with a sampling of days from Day −8 through Day 34. (A) Mouse BLI (flux), with a sampling of days from Day −8 through Day 34, for individual mice in Groups 1-3 in the mouse *in vivo* tumor control study.

**Supplementary Figure S11.**
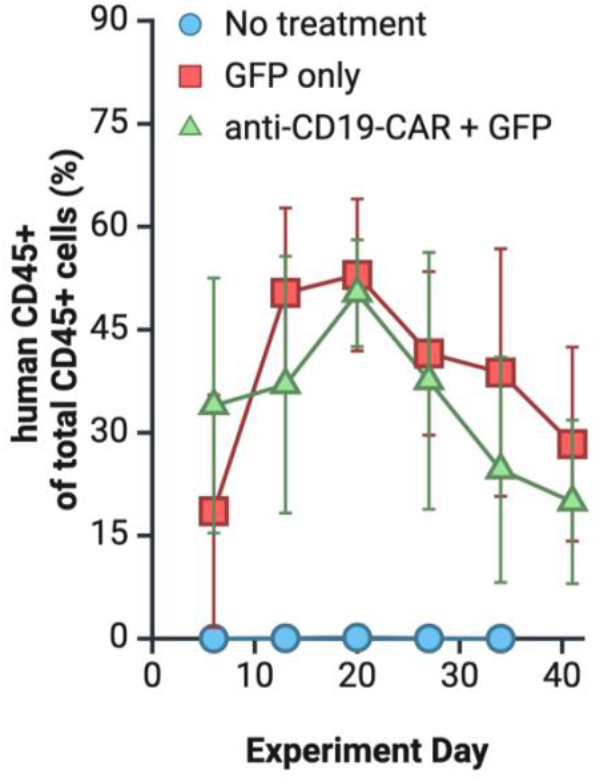
Human CD45+ cells as a percent of total CD45+ (human CD45+/[human CD45+ + mouse CD45+]) for Groups 2 and 3 of the *in vivo* mouse tumor control study, weekly measurements Day 6 through Day 34.

## SUPPLEMENTARY TABLES

**Supplementary Table S1.**

| Targeting Moiety | Surface Marker Positive Cells | Surface Marker Negative Cells |
| --- | --- | --- |
| CD3 | CD4+ and/or CD8+ | CD19+, CD20+ |
| CD4 | CD3+, TCRb+, CD8- | CD3+, TCRb+, CD8+ |
| CD8 | CD3+, TCRb+, CD4- | CD3+, TCRb+, CD4+ |
| CD19 | CD20+ | CD3+ |

**Supplementary Table S2.**
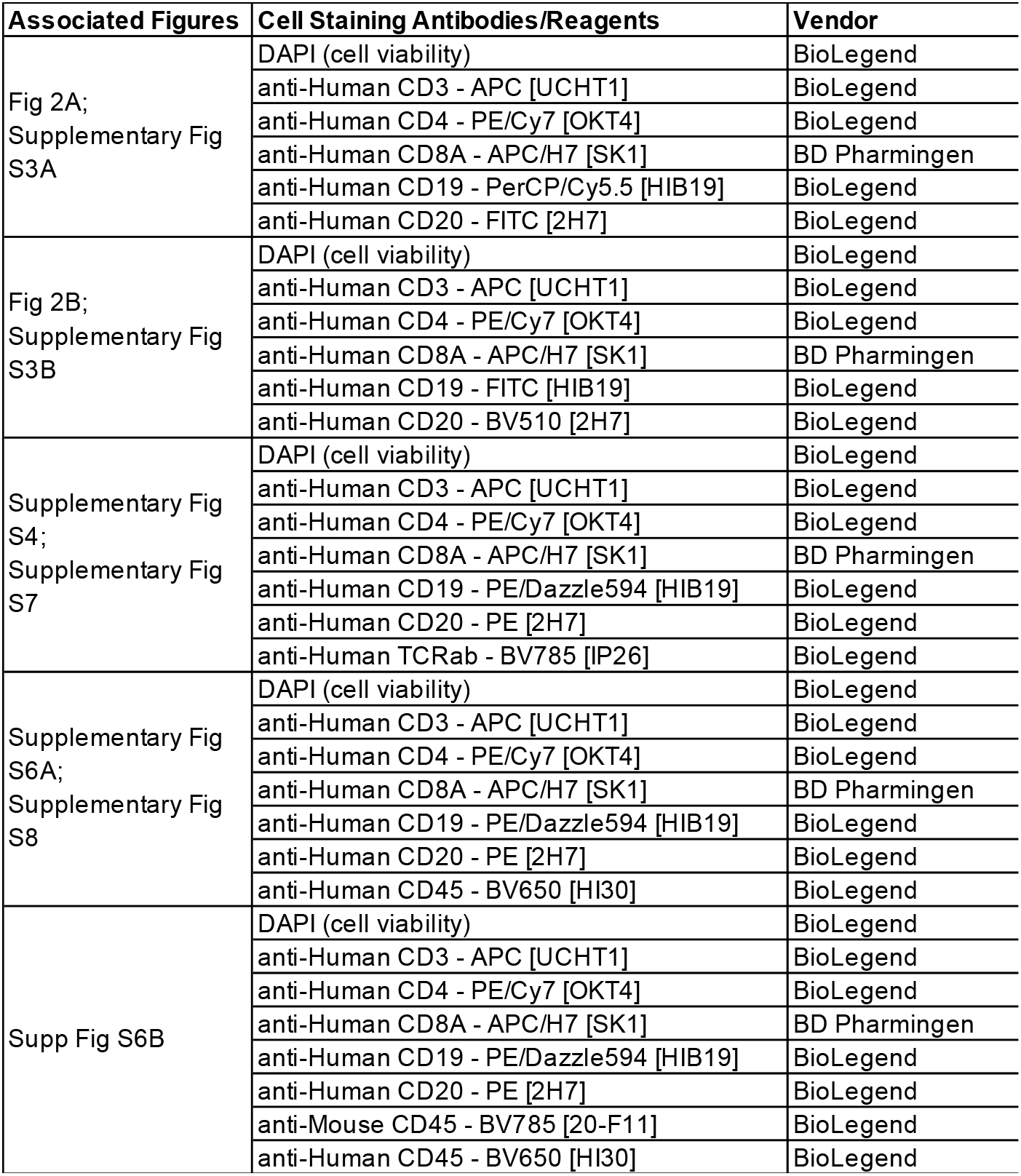

**Supplementary Table S3.**
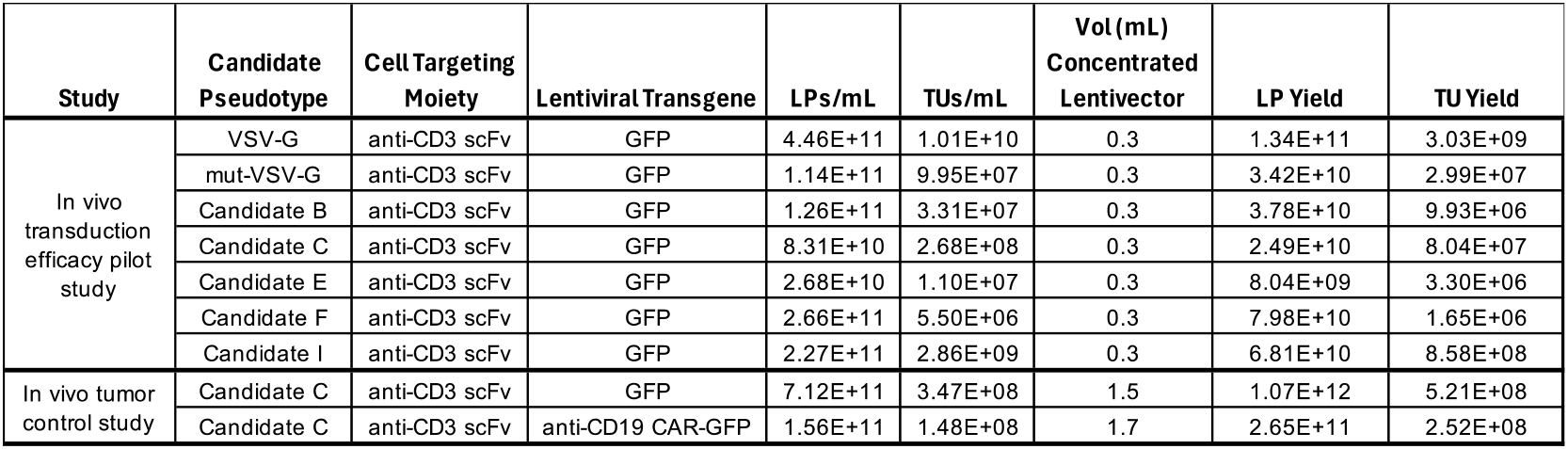

**Supplementary Table S4.**
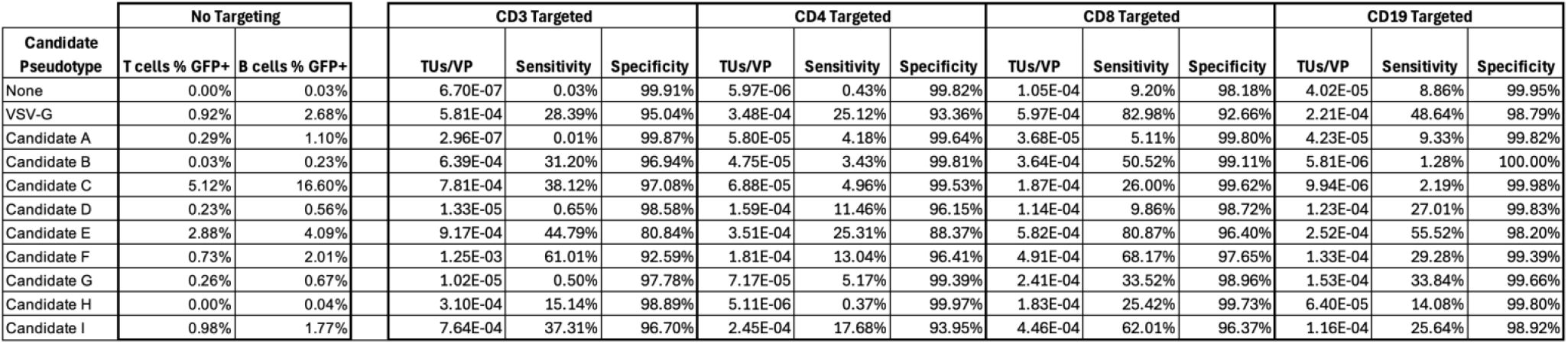

## Notes

### Competing Interest Statement

M.J. Spindler, A. Amezquita, E. Byrne, R. Edgar, S. Ravi1, S. Sandhu, T. Weller, and D.S. Johnson have received equity in GigaMune Inc. and/or cash salary for their work on the compositions and methods described in this manuscript.

